# A Novel Mutation in LXRα Uncovers a Role for Cholesterol Sensing in Limiting Metabolic Dysfunction-Associated Steatohepatitis (MASH)

**DOI:** 10.1101/2024.05.13.593869

**Authors:** Alexis T. Clark, Lillian Russo-Savage, Luke A. Ashton, Niki Haghshenas, Ira G. Schulman

## Abstract

Liver x receptor alpha (LXRα, Nr1h3) functions as an important intracellular cholesterol sensor that regulates fat and cholesterol metabolism at the transcriptional level in response to the direct binding of cholesterol derivatives. We have generated mice with a mutation in LXRα that reduces activity in response to endogenous cholesterol derived LXR ligands while still allowing transcriptional activation by synthetic agonists. The mutant LXRα functions as a dominant negative that shuts down cholesterol sensing. When fed a high fat, high cholesterol diet LXRα mutant mice rapidly develop pathologies associated with Metabolic Dysfunction-Associated Steatohepatitis (MASH) including ballooning hepatocytes, liver inflammation, and fibrosis. Strikingly LXRα mutant mice have decreased liver triglycerides but increased liver cholesterol. Therefore, MASH-like phenotypes can arise in the absence of large increases in triglycerides. Reengaging LXR signaling by treatment with synthetic agonist reverses MASH suggesting that LXRα normally functions to impede the development of liver disease.

## Introduction

Metabolic Dysfunction-Associated Steatotic Liver Disease (MASLD, formally referred to as non-alcoholic fatty liver disease) is estimated to impact approximately 25% of the adult population in the United Sates and 35% of adults worldwide ^1^. MASLD by itself can be relatively benign but approximately 25% of patients progress to Metabolic Dysfunction-Associated Steatohepatitis (MASH, formally referred to as non-alcoholic steatohepatitis), characterized by hepatocyte ballooning, inflammation of the liver, and fibrosis. Importantly, MASH increases the risk for cirrhosis, hepatocellular carcinoma, and liver failure ^2^. The processes triggering the progression of MASLD to MASH or even if there is a true stepwise progression from one pathological state to the other, however, remains to be determined. MASLD is characterized by excess fat accumulation in the liver and elevated rates of *de novo* fatty acid synthesis are often observed ^3, 4^. Since most MASLD patients do not have MASH, it is not clear if increases in liver fat alone drive the transition to MASH or if additional triggers are required.

In humans, liver cholesterol levels correlate with the severity of MASH ^5, 6, 7, 8^ and clinical studies indicate that patients with MASH can benefit from inhibiting cholesterol synthesis with statins ^9, 10, 11, 12, 13^. Furthermore, most high fat diet-dependent rodent models of MASH require also high levels of dietary cholesterol to promote disease ^14, 15, 16, 17^. High intracellular cholesterol is toxic leading to endoplasmic reticulum stress, inflammation, and cell death all of which have been suggested to contribute to MASH ^2^. Nevertheless, the mechanisms by which cholesterol directly influences the disease are not well established. The liver X receptors, LXRα (Nr1h3) and LXRβ (Nr1h2), are members of the nuclear hormone receptor superfamily of ligand activated transcription factors that function as important regulators of cholesterol and fatty acid metabolism. By directly binding cholesterol derivatives including oxidized forms of cholesterol (oxysterols) LXRs sense changes in intracellular cholesterol levels and regulate gene expression to maintain lipid homeostasis ^18, 19, 20^. Like many nuclear receptors, in the absence of agonist ligands LXRs can interact with corepressor proteins that inhibit transcription. Binding of either endogenous or synthetic agonists results in conformational changes in the LXR ligand binding domain that reduce interaction with corepressors and favor engagement with coactivators that increase transcription of LXR regulated genes ^21^. Genetic deletion of LXRα, the major LXR subtype expressed in the liver, impairs the catabolism and excretion of cholesterol and also decreases *de novo* fatty acid synthesis ^22, 23^. Consistent with the genetic data, molecular studies indicate that LXRs directly regulate the transcription of genes involved in cholesterol excretion, the synthesis of bile acids from cholesterol, and fatty acid synthesis ^22, 24, 25, 26^. Since elevated fatty acid synthesis is considered a major driver of steatosis in MASLD, inhibiting LXR activity has been put forward as a potential therapeutic approach for treating steatotic liver disease ^27^. Indeed, treatment with synthetic LXR antagonists reduces hepatic lipid accumulation in diet-dependent mouse models of liver steatosis ^28, 29, 30^. On the other hand, increasing LXR activity in mice with synthetic agonists promotes cholesterol excretion from the liver, the net movement of cholesterol out of the body, and reduces inflammation ^23, 31, 32^. Accordingly, even in the face of large increases in hepatic and plasma triglyceride levels, LXR activation has beneficial effects in models of chronic metabolic diseases including type II diabetes and atherosclerosis ^33, 34, 35, 36^. In support of the mouse models, genome wide association studies (GWAS) in humans have identified variants in the human LXRα gene (NR1H3) linked to lipid levels and insulin sensitivity ^37, 38, 39^. Rare variants in LXRα are also associated with markers of liver damage ^40^ suggesting an important role for LXRα in maintaining normal liver function.

We have generated mice with mutation of a conserved tryptophan at amino acid position 441 of the LXRα gene to phenylalanine (W441F). The W441F mutation impairs the transcriptional response of LXRα to endogenous cholesterol derived ligands while still allowing transcriptional regulation with potent synthetic agonists ^41, 42, 43^. LXRα W441F functions as a dominant negative receptor that inhibits LXR-dependent transcription *in vivo*. Mutant mice accumulate cholesterol in the liver and demonstrate increased pro-inflammatory and pro-fibrotic gene expression. In response to a high fat, high cholesterol diet W441F mice rapidly develop MASH-like phenotypes including ballooning hepatocytes, immune infiltration, and fibrosis. Reengaging LXR signaling by treating W441F mice with synthetic LXR agonists reverses cholesterol accumulation, inflammation, and fibrosis suggesting that LXRs normally function to impede the development of MASH.

## Results

### Characterization of LXRα W441F

Structural studies and reporter gene assays suggest that mutation of a conserved tryptophan in the carboxy-terminal helix 12 of the LXR ligand binding domain (amino acid 441 in mouse LXRα) to phenylalanine weakens pi-cation interactions with a histidine in helix 10 (amino acid 419 in mouse LXRα) that is required for the transcriptional response of LXRs to cholesterol derived ligands. Transcriptional activation by potent synthetic ligands such as T0901317, however, is less sensitive to the change to phenylalanine ^41, 42, 43^. To explore the effect of the W441F mutation in a more relevant setting, immortalized bone marrow derived macrophages derived from LXRα + LXRβ double knockout mice ^44^ were infected with adenovirus expressing mouse LXRα or LXRα W441F and the mRNAs encoding sterol regulatory element binding protein 1c (*Srebp1c*) and stearoyl CoA desaturase 1 (*Scd1*), two well characterized LXR target genes, were quantified. Cells infected with virus expressing GFP alone were used as controls (Figure 1A-B). As expected, wildtype LXRα increases gene expression in response to both the oxysterol 24(s),25-epoxycholesterol as well as the synthetic agonist T0901317. In contrast the response to 24(s),25-epoxycholesterol is selectively lost in cells expressing LXRα W441F. Dose response analysis indicates that the EC_50_ for T0901317 is increased by a factor of 5 in cells expressing LXRα W441F and that the maximum efficacy is reduced relative to cells expressing wildtype LXRα (Figure 1C). Western blotting indicates that the mutant protein is expressed at higher levels than the wildtype LXRα (Figure 1D-E) which is consistent with our published studies demonstrating that LXR half-life correlates with transcriptional activity ^45^.

**Figure 1.**
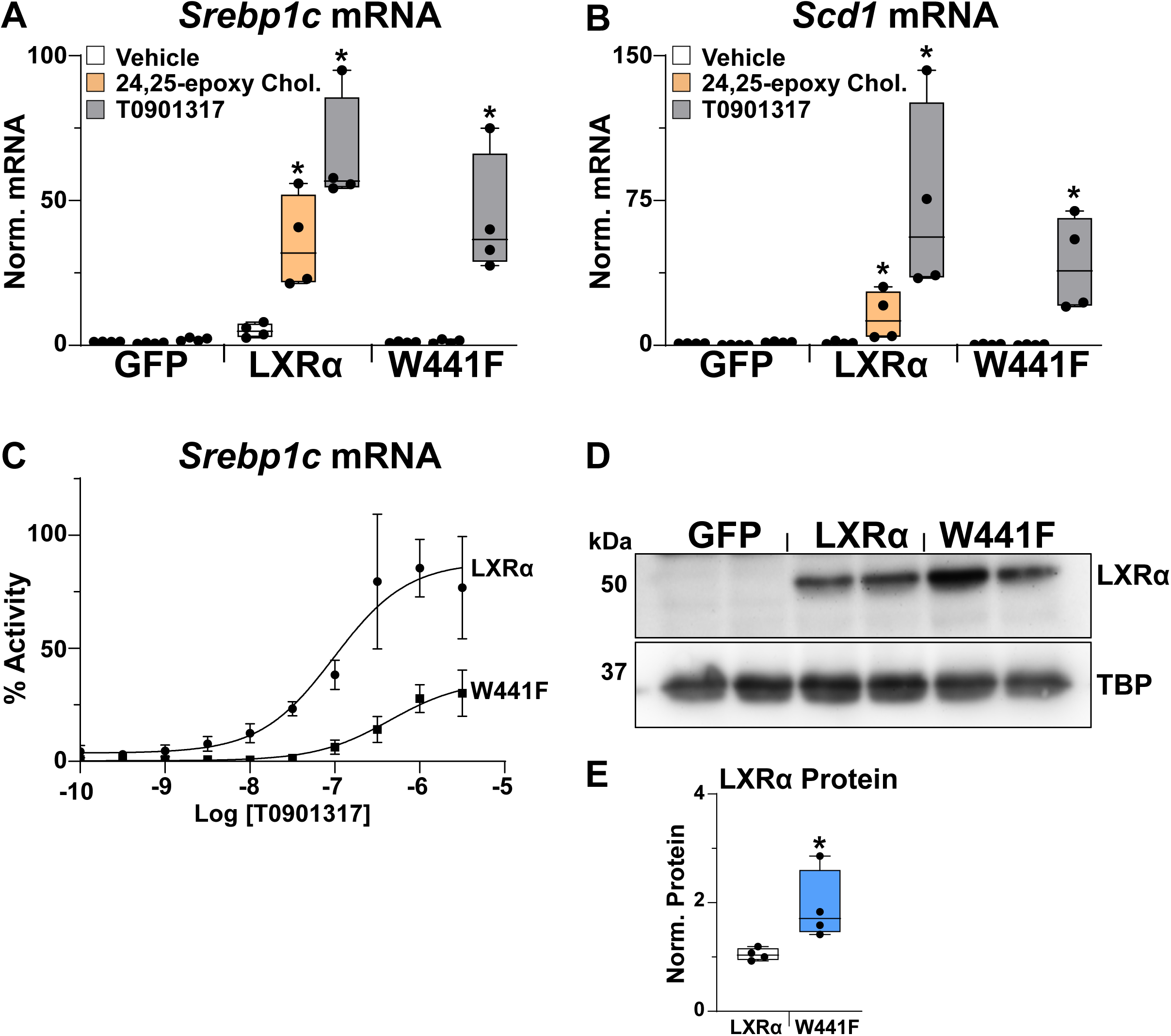
Characterization of LXRα W441F *in vitro*. Immortalized bone marrow derived macrophages were infected with adenovirus expressing LXRα or LXRα W441F. After infection, cells were treated for 24 hours with 10 µM 24(s),25-epoxycholesterol or 1.0 µM T0901317, RNA was isolated and the mRNA levels of **A**) *Srebp1c* and **B**) *Scd1* were measured by real-time PCR as described in the Methods section. *Statistically significant difference between vehicle and treated group for each virus determined by two-way ANOVA (*p* ≤ 0.05, n =4). The line in the middle of the bars represents the median. **C**) Cells were infected as described above and treated with different concentrations of T0901317 for 24 hours. RNA was isolated and *Srebp1c* mRNA levels were measured by real-time PCR as described in the Methods section. Data is the average and standard deviation of 3 independent experiments with *Srebp1c* mRNA measured in duplicate. **D**) Nuclear extracts were prepared from Immortalized bone marrow derived macrophages 24 hours after infection as described in the Methods. LXRα and TBP levels were examined by Western blotting. **E**) LXRα protein levels were quantified from Western by normalization to TBP using ImageQuant TL. Each point is an individual infection. The amount of wildtype LXRα was seat at 1. The line in the middle of the bars represents the median. *Statistically significant difference between LXRα and W441F determined by two-tailed Mann-Whitney test (*p* ≤ 0.05, n =4).

To examine the consequence of the W441F mutation *in vivo*, CRISPR was used to introduce the mutation into the *Lxrα* (*Nr1h3*) locus (Supplemental Figure 1A). Breeding of heterozygous mutant mice (referred to as at W/F) produces homozygous wildtype (referred to as W/W) and homozygous mutant (referred to as F/F) progeny at the expected mendelian ratios. At 10-12 weeks of age there are no significant differences in body weight among W/W, W/F and F/F females or males (Supplemental Figure 1B) and all mice appear normal. LXR regulated genes involved in cholesterol metabolism (Figure 2A-C) and fatty acid synthesis (Figure 2D-F), however, are significantly decreased in the livers and intestines of W441F mice compared to controls with relatively stronger decreases in homozygous (F/F) compared to heterozygous (W/F) mutant animals. Western blotting demonstrates approximately equal levels of LXRα protein in liver nuclear extracts from each of the 3 genotypes (Supplemental Figure 1D-E) indicating the observed changes in gene expression do not result from decreased W441F protein levels. There is a corresponding decrease in plasma triglycerides and cholesterol (Figure 2G-H) which reflects known roles for LXRs in promoting very low density lipoprotein (VLDL) secretion ^26, 46, 47^ and the production of high density lipoprotein (HDL) particles ^31, 48, 49^. Plasma glucose is slightly lower in mutant mice (Figure 2I), however, the decrease in glucose may be secondary to liver damage (see below). Consistent with the known role for LXRα in regulating bile acid synthesis ^22^, fecal bile acids are also decreased in W441F mice (Supplemental Figure 1C). Since LXRα W441F is not activated endogenous cholesterol derived ligands, we suggest that mutant receptors function as dominant negative inhibitors of transcription by binding to DNA but failing to activate transcription. Indeed, overexpressing LXRα W441F in AML12 mouse liver cells inhibits expression of LXR target genes (Supplemental Figure 1F-H).

**Figure 2.**
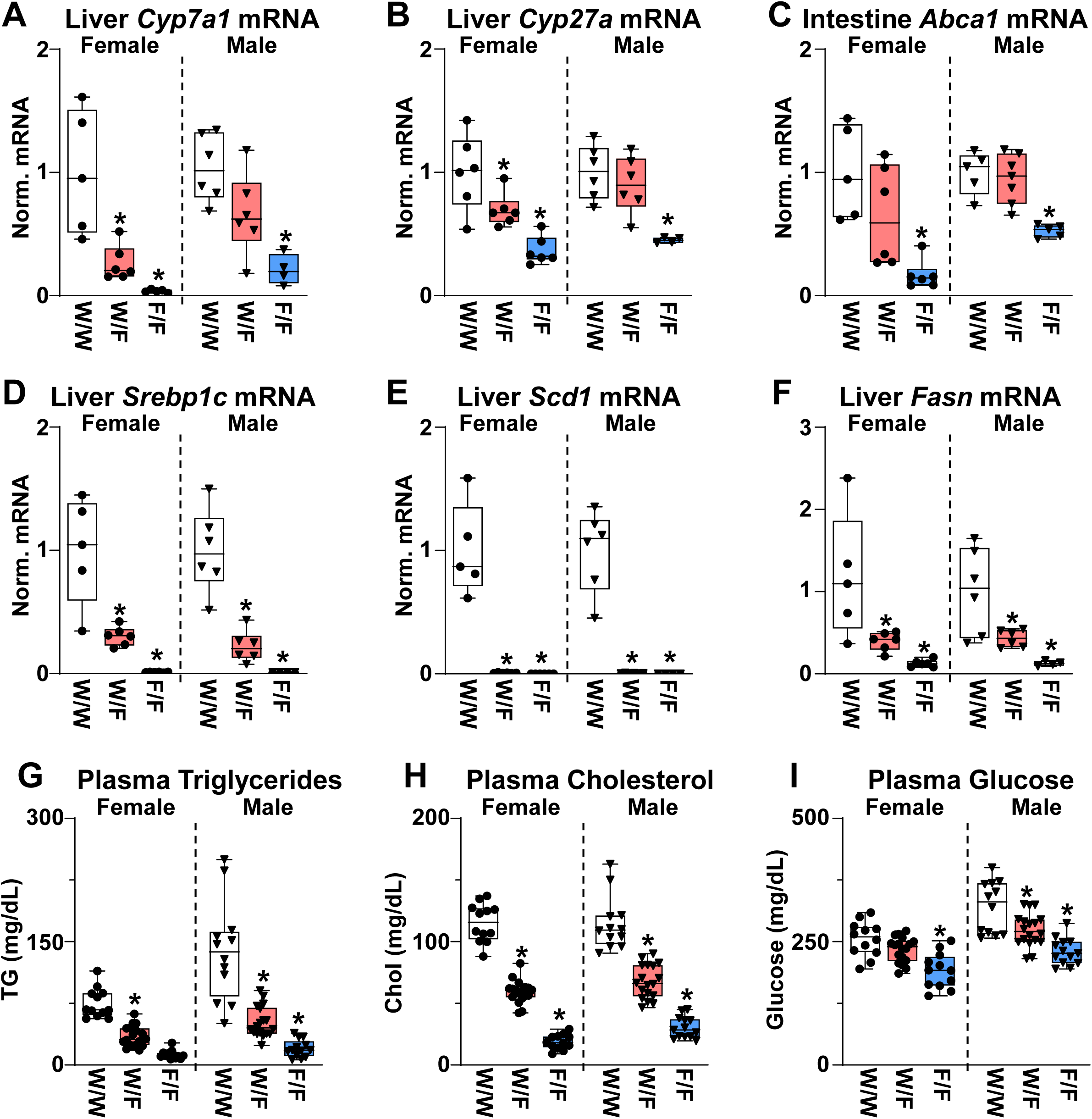
Gene expression and plasma lipids in LXRα W441F mice. Liver RNA, intestine RNA and plasma were isolated from 11-12 week old control (W/W), heterozygous mutant (W/F), and homozygous mutant (F/F) W441F mice maintained on normal chow diet. **A**-**F**) Gene expression measured by real-time PCR as described in the Methods section. *Statistically significant difference between control and mutant mice determined by one-way ANOVA (*p* ≤ 0.05, n = 4-6/group). **G**-**I**) Plasma triglycerides, cholesterol, and glucose were measured as described in the Methods section. *Statistically significant difference between control and mutant mice determined by one-way ANOVA (*p* ≤ 0.05, n = 12-18/group). The line in the middle of the bars represents the median.

Histological examination of liver sections by hematoxylin and eosin (H&E) staining uncovers evidence of immune infiltration in homozygous F/F W441F mice (Figure 3A). Additionally large pockets of neutral lipids are selectively detected by Oil Red staining of homozygous F/F mutant sections (Figure 3B). Quantification of tissue lipids indicates that liver triglycerides are decreased in both heterozygous W/F and homozygous F/F mice (Figure 3C). Liver cholesterol on the other hand is increased only in homozygous F/F mutant mice (Figure 3D-E) suggesting that the Oil Red O staining most likely identifies cholesterol loaded cells. The observation that LXR target genes involved in cholesterol catabolism are less affected in heterozygous mutants relative to genes involved in fatty acid synthesis (Figure 2A-F) may account for the gene dosage effects on hepatic cholesterol accumulation. Hepatomegaly, evidenced by larger liver to body weight ratios, is also observed selectively in F/F mice (Supplemental Figure 2A). Furthermore, increased staining with antibodies recognizing the proliferative marker Ki67 is detected in F/F hepatocyte nuclei (Figure 3F) indicating cell division is occurring. Previous studies have suggested that transcriptionally active LXRs constrain cell growth by limiting the availability of cholesterol needed to support proliferation ^50, 51, 52^. The increase in hepatocyte proliferation measured in homozygous F/F mutant mice supports the earlier work and further indicates that LXR signaling normally limits hepatocyte growth. Finally, the levels of alanine transaminase (ALT), and aspartate amino transferase (AST) are elevated in plasma from F/F mice, indicative of liver damage (Supplemental Figure 2B-C).

**Figure 3.**
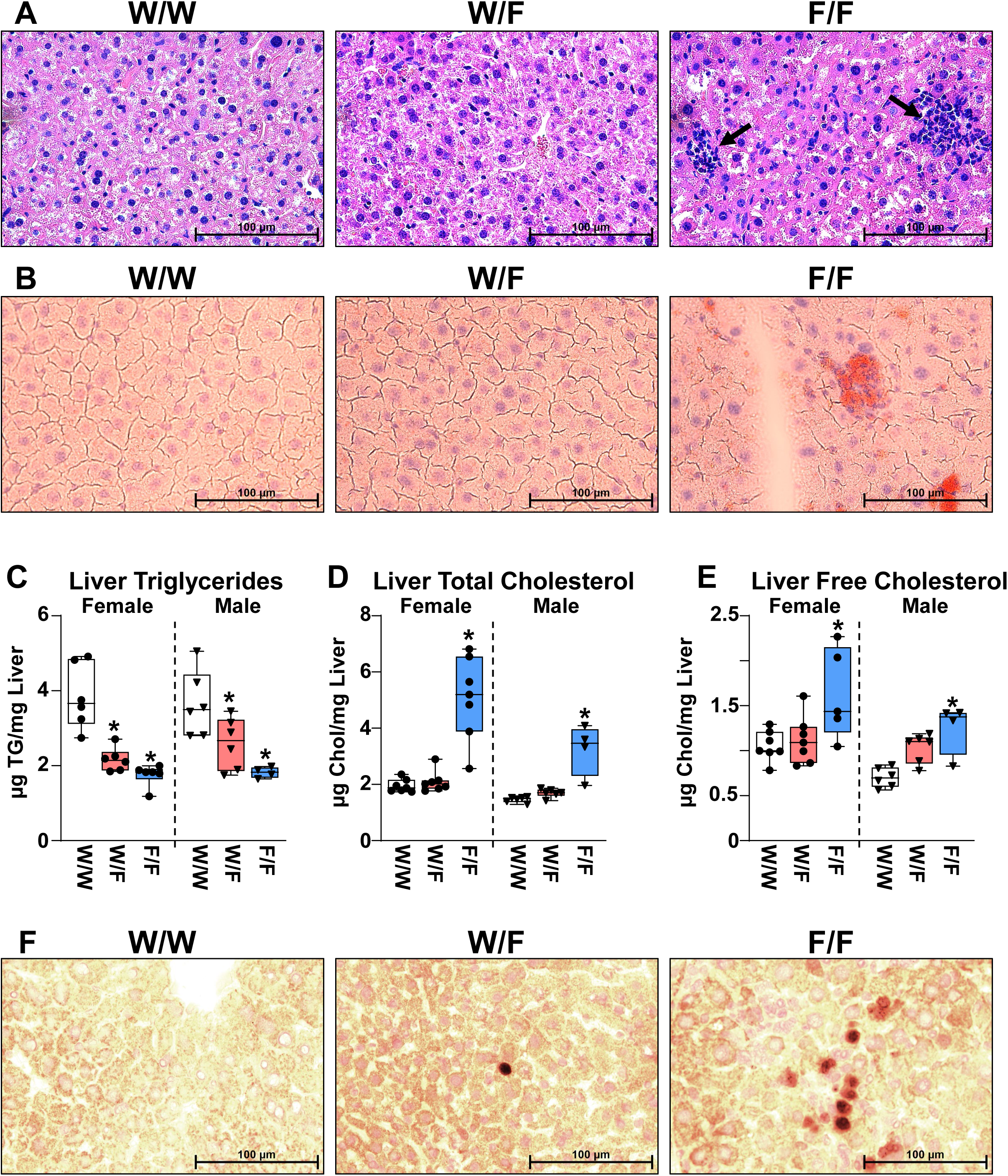
Cholesterol accumulation and increased cell division in LXRα W441F mice. **A**) H&E and **B**) Oil Red O staining of liver sections from representative male 11-12 week old control (W/W), heterozygous mutant (W/F), and homozygous mutant (F/F) W441F mice maintained on normal chow diet. Black arrows in **A** mark locations of immune infiltration. **C**-**E**) liver triglycerides and cholesterol were measured in samples from 11-12 week old control (W/W), heterozygous mutant (W/F), and homozygous mutant (F/F) W441F mice maintained on normal chow diet as described in Methods. *Statistically significant difference between control and mutant mice determined by one-way ANOVA (*p* ≤ 0.05, n = 4-6/group). The line in the middle of the bars represents the median. **F**) Ki67 staining of representative liver sections from 11-12 week old male control (W/W), heterozygous mutant (W/F), and homozygous mutant (F/F) W441F mice maintained on normal chow diet.

To further explore differences in hepatic gene expression among the 3 genotypes RNA-Sequencing (RNA-Seq) was used. Principle component analysis (PCA) of liver gene expression indicates that W/F heterozygotes more closely resemble W/W controls than heterozygous F/F mutants (Figure 4A). Compared to W/W controls, 59 genes are mis-regulated in heterozygous W/F livers (39 down and 20 up). Gene ontology analysis indicates that most of the down-regulated genes are linked to fatty acid metabolism (Figure 4B) and 18 are either well established LXR target genes or genes that have nearby LXR binding sites identified by chromatin immunoprecipitation sequencing ^53^. The only significant up-regulated pathways in W/F mice include 4 genes induced 2-2.3 fold that are associated with sterol metabolism. In contrast, over 2000 genes are differentially expressed in F/F homozygous mutant livers compared to W/W controls (488 down and 1688 up). Down regulated genes in F/F are also largely linked to fatty acid metabolism. Major gene networks selectively up regulated in F/F livers include those associated with cell division, inflammation/immune cell function, and cell adhesion/extracellular matrix (Figure 4B). Representative changes in gene expression were confirmed by quantitative PCR (Figure 4C-H). Increased expression of the chemokine receptor *Cx3cr1* (Figure 4C) along with elevated expression of pro-inflammatory genes such as *Tnf* (Figure 4D) is consistent with the immune infiltration observed by H&E staining and suggests a pro-inflammatory environment in homozygous F/F mutant livers. Immunohistochemical staining with antibodies recognizing the macrophage marker Cd68 also detects increased numbers of macrophage in F/F mutant livers (Figure 4I). Furthermore, several genes reported to be markers of lipid associated macrophage populations defined in models of MASH and other chronic disease settings including *Trem2*, and *Spp1* ^54, 55, 56^ are increased in F/F livers (Figure 4E-F) while expression of Kupffer cell markers such as Clec4f ^57, 58, 59, 60^ are decreased (Figure 4I). The decrease in Kupffer cell markers detected in homozygous F/F mutant mice is supported by previous studies identifying LXRα as a lineage determining factor for this cell type ^57, 58, 59^ and suggests sterol-dependent signaling is required for this function. Similar differences in gene expression are observed when RNA is isolated from an enriched population of liver macrophages (Supplemental Figure 3A-D). The changes in macrophage gene expression suggest that LXR signaling is required for the proper specification and function of liver myeloid populations. By phagocytosing dead and dying cells throughout the body, macrophages are required to respond to large increases in intracellular cholesterol levels. We observe significant enlargement of spleens in homozygous F/F mice which are characterized by the appearance of lipid loaded macrophages as determined by co-localization of Oil Red O staining with the macrophage marker Cd68 (Supplemental Figure 3E-G). Thus, W441F mice uncover important roles for cholesterol sensing by LXRs in liver and splenic macrophage populations.

**Figure 4.**
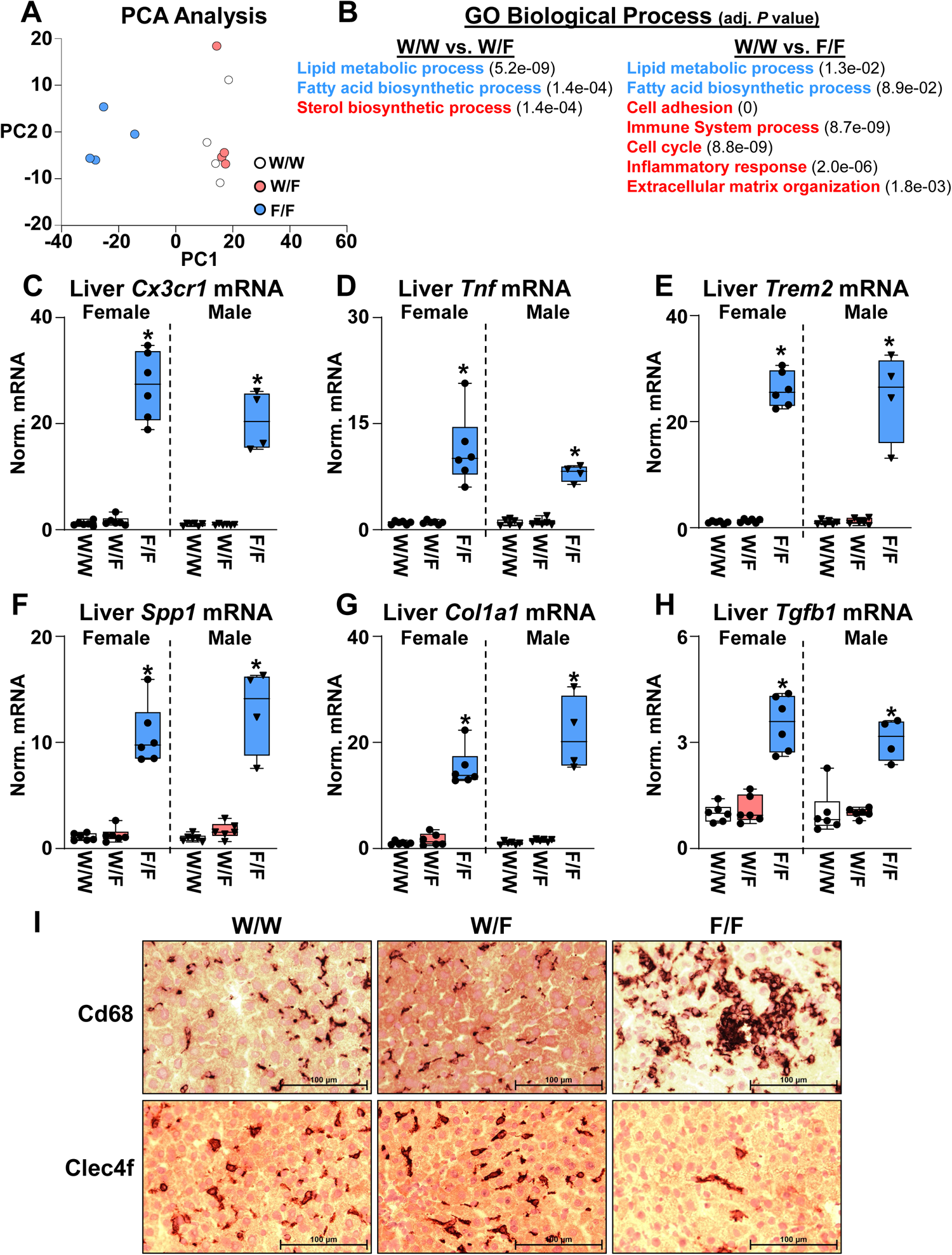
Liver gene expression in LXRα W441F mice. RNA-Seq was carried out as described in the Methods section using liver RNA isolated from 11-12 week old male control (W/W), heterozygous mutant (W/F), and homozygous mutant (F/F) W441F mice maintained on normal chow diet (n=4 group). **A**) PCA analysis. **B**) Summary of enriched GO terms in pairwise comparisons. **C**-**H**) Gene expression measured by real-time PCR as described in the Methods section. *Statistically significant difference between control and mutant mice determined by one-way ANOVA (*p* ≤ 0.05, n = 4-6/group). The line in the middle of the bars represents the median. **I**) Cd68 (top) and Clec4f (bottom) staining of representative liver sections from 11-12 week old female control (W/W), heterozygous mutant (W/F), and homozygous mutant (F/F) W441F mice maintained on normal chow diet.

### Rapid Development of MASH Phenotypes

On the normal mouse diet heterozygous F/F mutant livers express many phenotypes associated with MASH including lipid loaded hepatocytes, immune infiltration, and pro-fibrotic gene expression (e,g. *Spp1*, *Col1a1*, *Tgfb1*. Figure 4F-H). Nevertheless, deposition of extracellular matrix and histological evidence of fibrosis is not detected. To examine the requirement for LXR signaling under MASH promoting conditions, 11-week-old W441F mice were fed a MASH promoting diet (21% fat, 1.25% cholesterol, 34% sucrose) for 2 weeks. Representatives of each genotype were sacrificed at this time point while the remaining animals continued on diet and were treated with vehicle or the LXR agonist T0901317 for an additional 2 weeks to reactivate LXR signaling (total of 4 weeks exposure to diet). After 2 weeks on diet there is evidence of collagen deposition detected by Picrosirius Red staining in both heterozygous and homozygous W441F mice that further increases at 4 weeks in vehicle treated mice (Figure 5A-C). Since most high fat/high cholesterol diet-dependent MASH models take several months to develop histologically detectable fibrosis ^15, 16, 17^ the rapid development of fibrosis in W441F mice points to a critical role for LXR activity in regulating this process.

**Figure 5.**
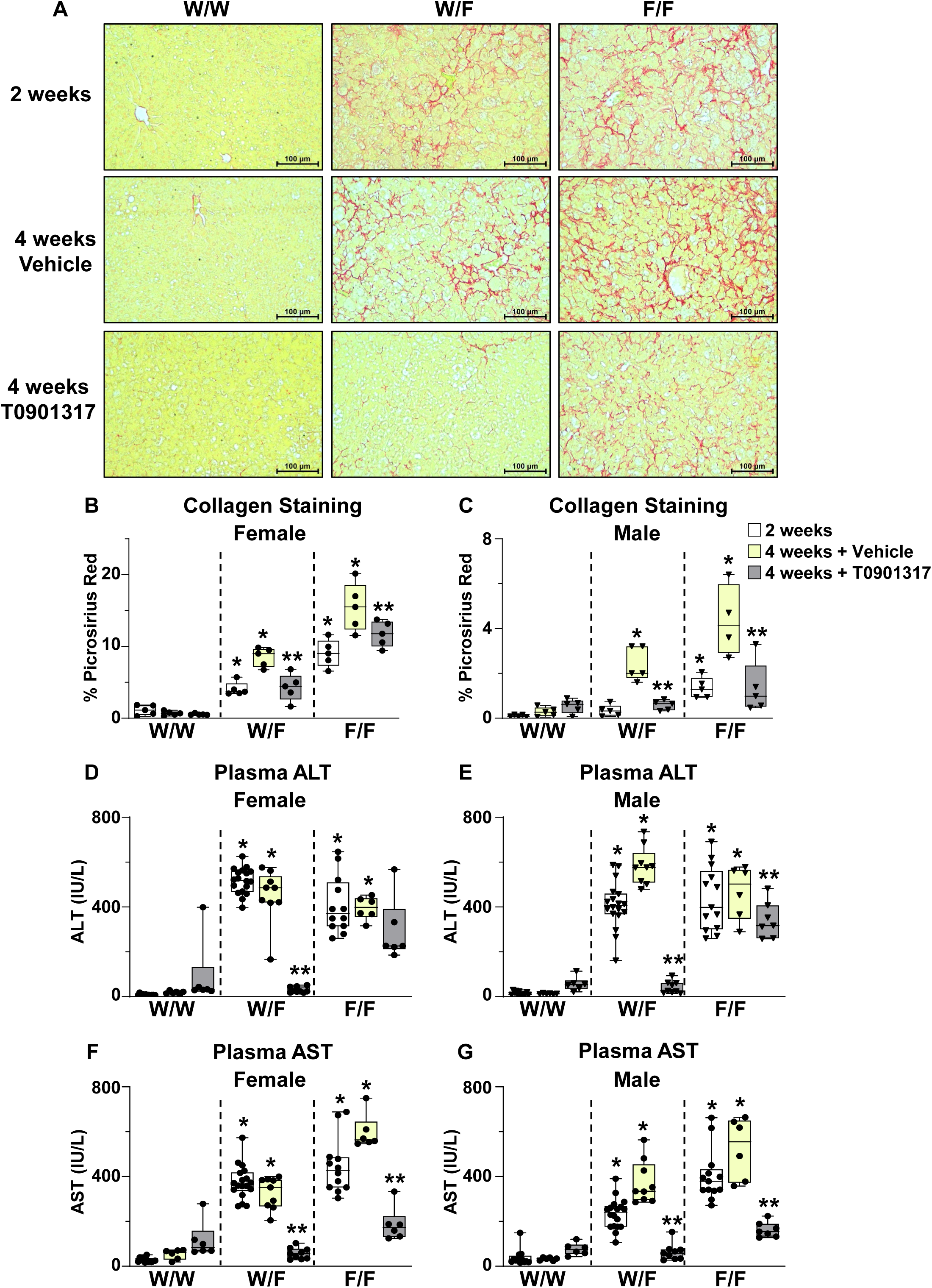
Fibrosis and liver damage in LXRα W441F mice. Female and male 11-12 week old control (W/W), heterozygous mutant (W/F), and homozygous mutant (F/F) W441F mice were fed the MASH diet. A subset of mice was sacrificed after 2 weeks. The remaining mice were maintained on the MASH diet for an additional 2 weeks and treated with vehicle or 10 mg/kg T0901317 daily by oral gavage as described in the Methods section. **A**) Liver sections from representative female mice stained with Picrosirius Red to visualize collagen. **B**-**C**) Quantification of the % surface area positive for Picrosirius Red in liver sections from female and male mice as described in the Methods section. *Statistically significant difference between control and mutant mice in the same treatment group; **statistically significant difference between vehicle and T0901317 treated mice of the same genotype determined by two-way ANOVA (*p* ≤ 0.05, n = 5/group). **D**-**G**) Plasma ALT and AST levels. *Statistically significant difference between control and mutant mice in the same treatment group; **statistically significant difference between vehicle and T0901317 treated mice of the same genotype determined by two-way ANOVA (*p* ≤ 0.05, n = 6-12/group). The line in the middle of the bars represents the median.

Strikingly, collagen staining is reduced after reactivation of LXR signaling by T0901317 compared to vehicle treated animals and resembles the levels measured in animals at 2 weeks. Liver damage quantified by plasma ALT and AST activity follows a similar trajectory and is also strongly reduced by reengaging LXR activity (Figure 5D-G). Notably liver enzymes are normalized in heterozygous (W/F) mutant mice.

Oil Red O staining of neutral lipids in liver sections from mice on the MASH diet (Figure 6A) illustrates unique patterns of lipid accumulation in W441F mice compared to controls. As typically observed in livers from mice fed high fat diets, control W/W mice have relatively small intensely stained lipid droplets that increase in size between 2 and 4 weeks. Furthermore, treatment of control mice with the LXR agonist T0901317, known to promote fatty acid synthesis ^25, 26^, increases the size of the droplets. In contrast, hepatocytes from heterozygous and homozygous W441F mice exhibit diffuse lipid staining (Figure 6A). The lipid loaded cells in W441F mice appear to correlate with large ballooning hepatocytes detected in H&E stained liver sections (Figure 6B) and we suggest that these cells are filled largely with cholesterol. Consistent with this hypothesis, liver cholesterol is elevated in W441F mice after 2 and 4 weeks of diet while liver triglycerides are decreased compared to W/W controls (Figure 6C-F). Reactivation of LXR signaling by treatment with T0901317 reduces liver cholesterol in heterozygous W/F mutants but does not drive a significant reduction in homozygous F/F mice (Figure 6). Livers from MASH diet treated W441F mice are extremely fibrotic (Supplemental Figure 4A) and difficult to extract such that the amount of cholesterol and the effect of reactivating LXR may be underestimated particularly in F/F samples. Liver and spleen weights are also significantly increased in W441F mice after 2 and 4 weeks on the MASH diet and re-engaging LXR signaling by treatment with T0901317 either completely (W/F heterozygotes) or partially (F/F homozygous) rescues these phenotypes (Supplemental Figure 4B-E). The increase in liver weight observed in W/W control mice treated with T0901317 is consistent with previous reports of this compound promoting hepatomegaly in high fat fed mice ^26^. Finally, W441F mutant mice fail to gain weight on the MASH diet and their fat mass is reduced (Supplemental Figure 5). This failure to thrive appears to be secondary to liver damage and is reversed in T0901317 treated animals.

**Figure 6.**
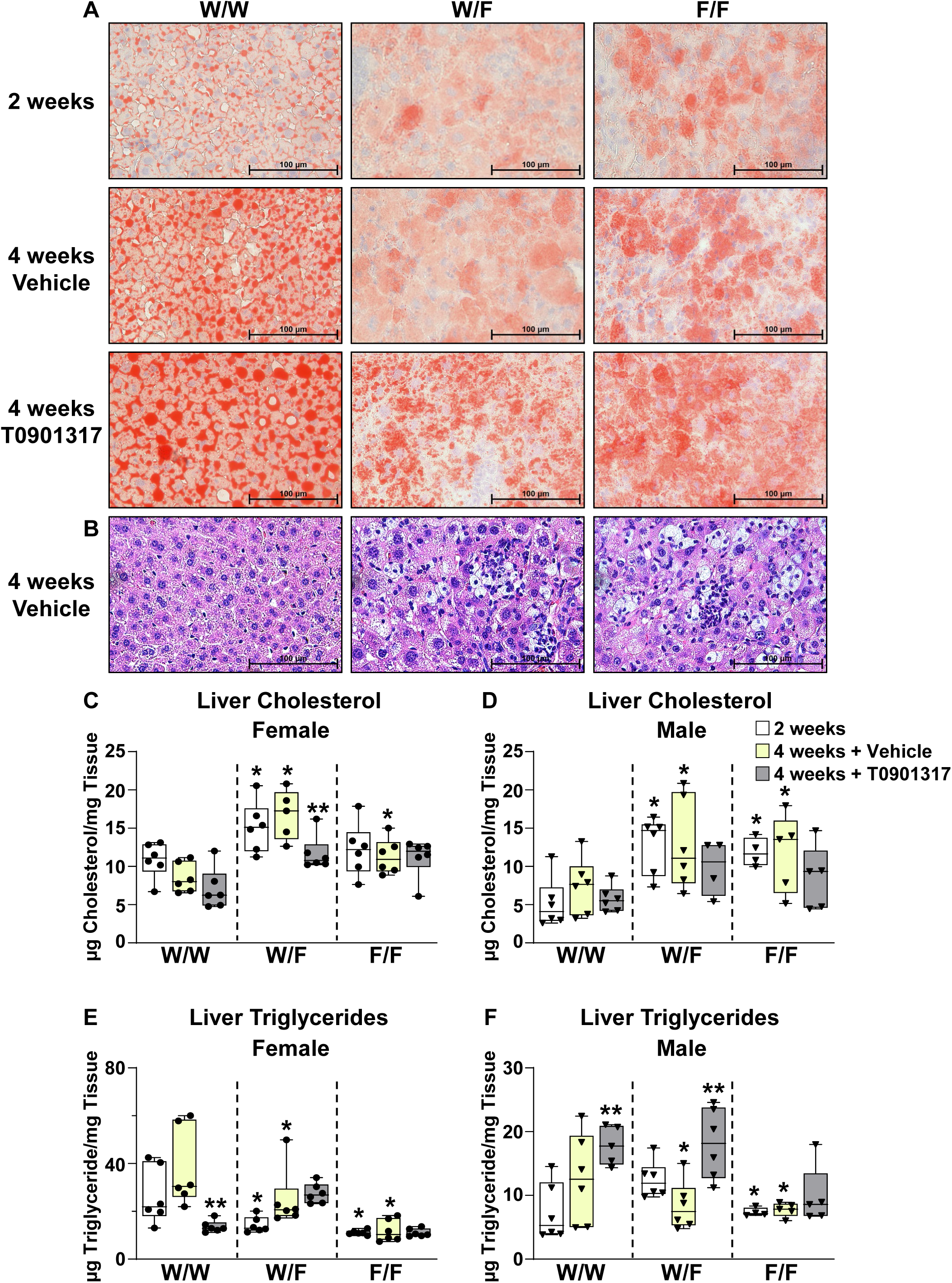
Lipid accumulation in livers from LXRα W441F mice fed the MASH diet. Female and male 11-12 week old control (W/W), heterozygous mutant (W/F), and homozygous mutant (F/F) W441F mice were fed the MASH diet. A subset of mice was sacrificed after 2 weeks. The remaining mice were maintained on the MASH diet for an additional 2 weeks and treated with vehicle or 10 mg/kg T0901317 daily by oral gavage as described in the Methods section. Representative liver sections from female mice stained with **A**) Oil Red O and **B**) H&E. **C**-**F**) Liver cholesterol and triglyceride levels measured as described in the Methods section. *Statistically significant difference between control and mutant mice in the same treatment group; **statistically significant difference between vehicle and T0901317 treated mice of the same genotype determined by two-way ANOVA (*p* ≤ 0.05, n = 5/group). The line in the middle of the bars represents the median.

After 2 weeks on the MASH diet W/F heterozygote livers now phenotypically resemble F/F homozygous livers demonstrating increased hepatic cholesterol, immune infiltration, and collagen deposition. Changes in gene expression are an additional manifestation of the influence of diet on the heterozygous W/F phenotype. After 4 weeks on the MASH diet, PCA of liver RNA-Seq data indicates that liver gene expression in vehicle treated W/F heterozygous mice now closely resembles the expression pattern observed in F/F homozygotes with 4030 and 5095 mis-regulated genes in W/F and F/F livers respectively (Figure 7A). GO analysis identifies many of the same pathways seen mis-regulated in normal chow F/F mice including lipid metabolism, inflammation, and cell adhesion/extracellular matrix, however the magnitude of the changes in gene expression are greater after exposure to the MASH diet (Figure 7B-J). Heterozygous W/F mutant mice are particularly responsive to T0901317 treatment and PCA of liver RNA-Seq data sets indicate that the transcriptome of LXR agonist treated W/F heterozygotes closely mimics the transcriptome of vehicle treated control W/W mice (Figure 7A). The diet-dependent changes in liver gene expression measured in W/F heterozygotes uncovers a context selective haploinsufficiency of the W441F mutation. Strikingly, in mutant mice treated with T0901317 to reactivate LXR the MASH phenotypes including lipid accumulation, immune infiltration and fibrosis measured at the level of gene expression or by histology are largely reversed (Figure 5-7).

**Figure 7.**
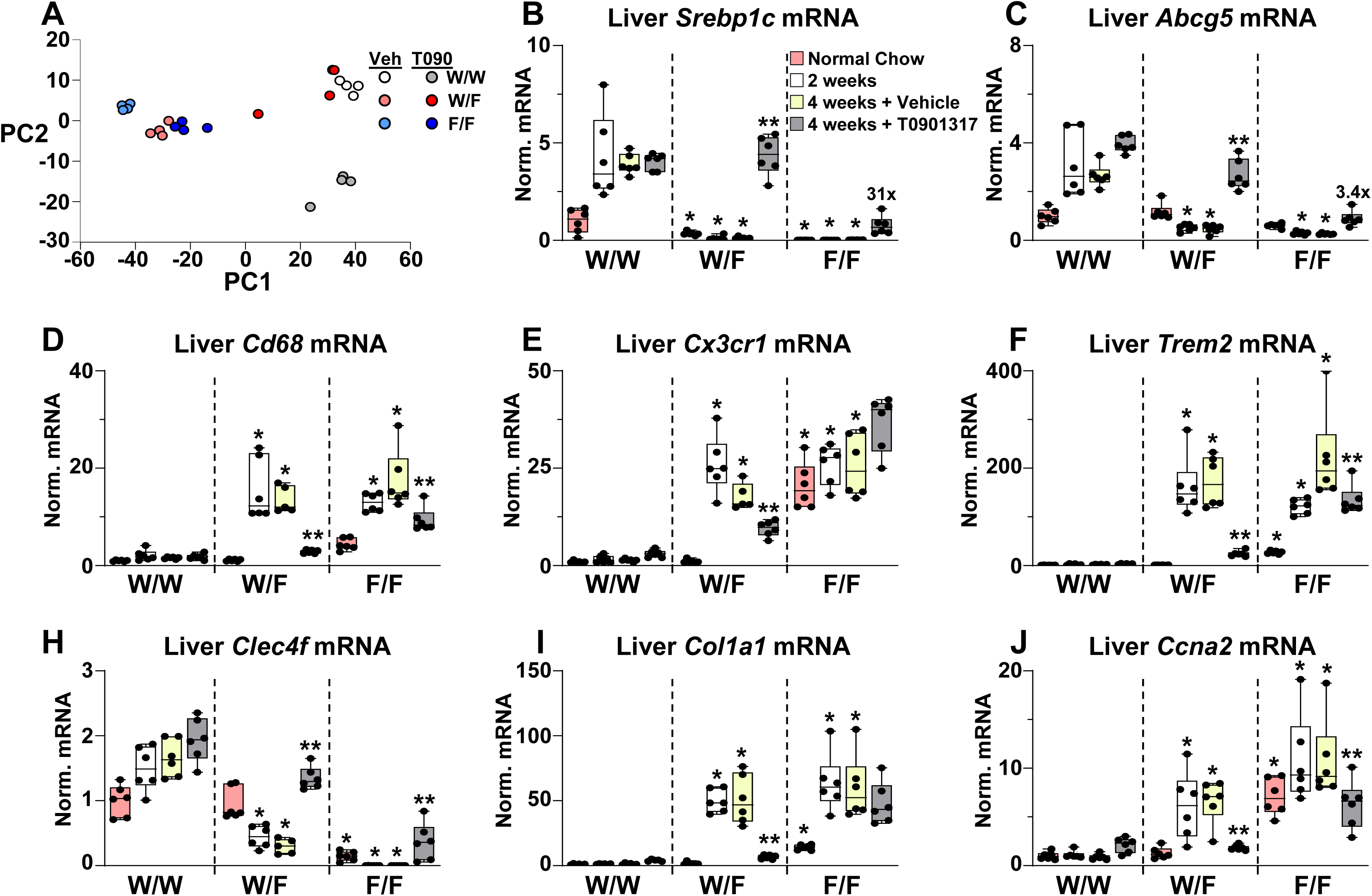
Liver gene expression in LXRα W441F mice fed the MASH diet. Female and male 11-12 week old control (W/W), heterozygous mutant (W/F), and homozygous mutant (F/F) W441F mice were fed the MASH diet. A subset of mice was sacrificed after 2 weeks. The remaining mice were maintained on the MASH diet for an additional 2 weeks and treated with vehicle or 10 mg/kg T0901317 daily by oral gavage as described in the Methods section. **A**) PCA analysis of RNA-Seq data using liver RNA isolated from male mice (n = 4/group). **B**-**J)** Gene expression in livers from female mice measured by real-time PCR as described in the Methods section. Gene expression was also measured in an independent cohort of 11-12 week old female mice maintained on normal chow. Numbers in **B** and **C** represent the fold induction by T0901317 in F/F mice. *Statistically significant difference between control and mutant mice in the same treatment group; **statistically significant difference between vehicle and T0901317 treated mice of the same genotype determined by two-way ANOVA (*p* ≤ 0.05, n = 6/group). The line in the middle of the bars represents the median.

TAZ is a transcriptional coactivator that is over expressed in human MASH patients and in mouse models ^61, 62, 63^. Recent studies indicate that TAZ is stabilized by elevated hepatic cholesterol and promotes fibrosis and inflammation ^63, 64, 65^. We do not detect differences in TAZ protein levels in livers from female control, heterozygous or homozygous W441F mice fed the MASH Diet (Supplemental Figure 6A-B). In contrast, there is an increase in TAZ levels in livers from male W441F mice (Supplemental Figure 6C-D). Interestingly, the published studies exploring the contribution of TAZ to the development of MASH in mice have only examined male animals ^63, 64, 65^. Treatment of male mice with T0901317 for 2 weeks increases the amount of TAZ in control W/W mice (Supplemental Figure 6C-D). Nevertheless, there is no histological evidence of inflammation or fibrosis in these animals. Therefore, in the W441F model increasing TAZ is not sufficient to promote MASH at least after 4 weeks of exposure to a MASH diet.

### Hepatocyte-Specific Expression of W441F Drives MASH

LXRα is expressed in all the cell-types that contribute to MASH including hepatocytes, Kupffer cells, infiltrating macrophages, and hepatic stellate cells ^19^. Aberrant lipid accumulation in hepatocytes, however, is thought to be the original insult driving MASH in humans and in mouse models ^2, 3, 66^. To determine if expression of LXRα W441F selectively in hepatocytes is sufficient to promote MASH phenotypes in mice we used CRISPR to integrate a trans-gene expressing FLAG tagged LXRα W441F under control of a Cre-dependent Lox-Stop-Lox cassette at the *Rosa26* locus. The *Rosa26* insertion was crossed into *Lxra*^fl/fl^ mice ^23^ such that Cre promotes expression of LXRα W441F while simultaneously excising the wildtype allele. To selectively express W441F in hepatocytes *Rosa26* W441F + *Lxrα*^fl/fl^ mice were infected with adeno associated virus (AAV) expressing GFP or Cre under control of the hepatocyte-specific thyroxine binding globulin (TBG) promoter (Figure 8A). Following infection, mice were maintained on a normal chow diet for 2 weeks before being switched to the MASH diet for an additional 4 weeks. Western blotting of liver nuclear extracts indicates that mice infected with TBG-Cre express approximately 10 times more FLAG-W441F protein than TBG-GFP controls (Figure 8B and Supplemental Figure 7A). After 4 weeks on the MASH diet, mice expressing W441F in hepatocytes demonstrate essentially the same phenotypes observed in mice expressing W441F from the endogenous locus including lipid loaded ballooning hepatocytes (Figure 8C-D), decreased liver triglycerides (Figure 8E), increased liver cholesterol (Figure 8F), diffuse lipid staining (Figure 8G-H), increased collagen deposition (Figure 8I-K), elevated liver enzymes (Figure 8L), and enlarged livers (Supplemental Figure 7B). Hepatocyte-selective W441F mice also fail to gain weight (Supplemental Figure 7C-D) indicating that this phenotype is autonomous to hepatocytes and is most likely a secondary effect of liver damage. Direct LXR target genes such as *Srebp1c* and *Abcg5* are down-regulated (Figure 8M-N). Conversely markers of infiltrating macrophages, lipid associated macrophages, fibrosis, and cell division are up-regulated (Figure 8O-T). Changes in Cre-dependent changes in gene expression are not observed in the intestine (Supplemental Figure 7E-G). The characterization of hepatocyte selective W441F mice indicates that inhibiting LXR transcriptional activity in hepatocytes is sufficient to rapidly promote phenotypes associated with MASH.

**Figure 8.**
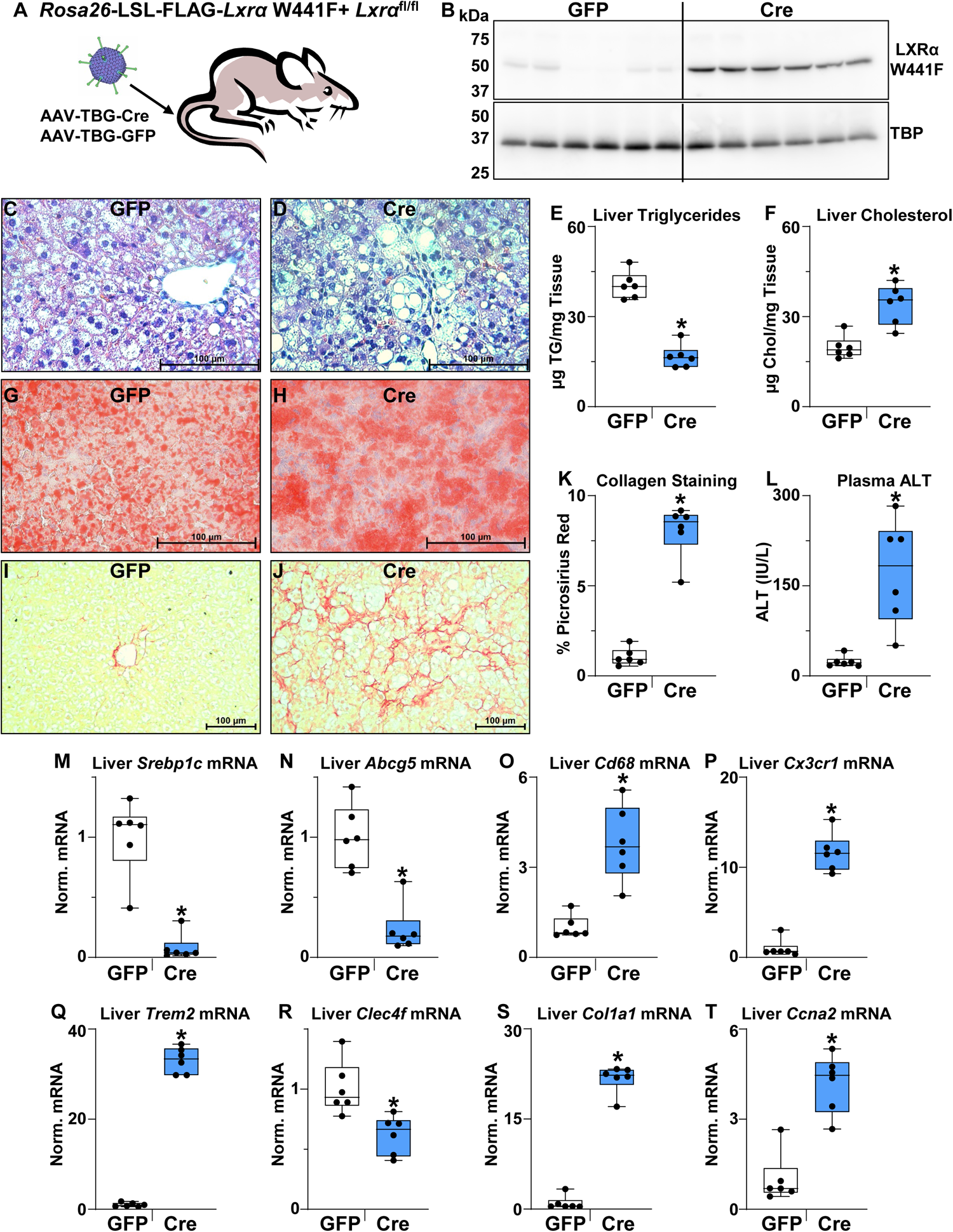
Hepatocyte-selective expression of LXRα W441F promotes MASH. Female 9 week old *Rosa26*-LSL-FLAG-LXRα W441F/*Lxrα*^fl/fl^ mice were infected with AAV-TBG-GFP or AAV-TBG-Cre viruses. Mice were maintained on a normal chow diet for 2 weeks and then fed the MASH diet for 4 weeks. **A**) Experimental schematic. **B**) Liver nuclear extracts were prepared as described in the Methods. FLAG-LXRα and TBP levels were examined by Western blotting. **C**-**D**) Representative H&E stained liver sections. **E**-**F**). Liver triglycerides and cholesterol were measured as described in the methods section. *Statistically significant difference between GFP and Cre mice determined by unpaired two-tailed t-test (*p* ≤ 0.05, n = 6/group). **G**-**H**) Representative Oil Red O stained liver sections. **I**-**J**) Representative Picrosirius Red stained liver sections. **K**) Quantification of the % surface area positive for Picrosirius Red in liver sections as described in the Method section. **L**) Plasma ALT measured as described in the Methods section. *Statistically significant difference between GFP and Cre mice determined by two-tailed Mann-Whitney tests (*p* ≤ 0.05, n = 6/group). **M-T**) Liver gene expression measured by real-time PCR as described in the Methods section. *Statistically significant difference between GFP and Cre mice determined by unpaired two-tailed t-test or two-tailed Mann-Whitney test (*p* ≤ 0.05, n = 6/group). The line in the middle of the bars represents the median.

## Discussion

MASH is rapidly becoming the leading cause of liver disease, liver failure and hepatocellular carcinoma in humans ^1^. Although steatotic liver disease (fatty liver) is reaching epidemic proportions due to the rise in obesity ^1^, what drives the transition of steatosis to the inflammation and fibrosis characteristic of MASH is not known. LXRα is an intracellular cholesterol sensor that regulates lipid metabolism at the transcriptional level in response to the direct binding of cholesterol derivatives ^19^. We demonstrate that disrupting the ability of LXRα to sense cholesterol generates a dominant negative transcription factor (W441F) that inhibits LXR dependent transcription even when present in a heterozygous state. Within 2 weeks of exposure to a high fat, high cholesterol MASH promoting diet W441F mice develop the hallmarks of human MASH including lipid laden ballooning hepatocytes, elevated liver enzymes, inflammation, and fibrosis. W441F impairs the transcriptional response to endogenous cholesterol derived ligands, however, the mutant receptor can still activate transcription in response to potent synthetic agonists such as T0901317. Importantly, reactivation of LXR signaling after 2 weeks of the MASH diet blocks the further progression of fibrosis and either completely (heterozygotes) or partially (homozygotes) reverses liver damage and inflammation. W441F mice accumulate high levels of hepatic cholesterol while hepatic triglycerides levels are decreased highlighting the potential pathological role of cholesterol in MASH. The cholesterol associated steatohepatitis (CASH) observed in W441F mice is consistent with studies in humans linking the severity of MASH with hepatic cholesterol levels ^5, 6, 8^. LXRα also is a transcriptional regulator of fatty acid synthesis and inhibition of LXR activity with synthetic antagonists has been suggested as a therapeutic approach for treating steatotic liver disease ^27^. Although it is often difficult to extrapolate genetic manipulations to the pharmacological action of small molecules, our studies suggest that agents such as LXR antagonists that could increase hepatic cholesterol levels may not be beneficial for patients withpre-existing MASH.

Studies of MASH in mouse models have described an influx of Cx3cr1 expressing myeloid cells that appear to give rise to lipid associated macrophages expressing high levels of Trem2 and Gpnmb ^54, 55, 56^. Similar populations of lipid associated macrophages have been detected in models of other chronic diseases associated with high cholesterol levels including atherosclerosis, diabetes, and Alzheimer’s disease ^67, 68, 69^. The functions of lipid associated macrophages and if they have beneficial or detrimental effects in different pathological settings is an area of active inquiry. In MASH there is also a loss of tissue resident macrophages (Kupffer cells) that has been attributed to a loss of identity and/or cell death ^57, 58, 59, 60^. Similar increases in lipid associated macrophages and decreases in Kupffer cell markers are detected in W441F mice. The decrease in Kupffer cells detected in W441F mice is consistent with the known role for LXRα as a lineage determining factor for these cells ^56, 57, 58, 59^. Strikingly, the macrophage landscape in W441F mice rapidly reverts to control condition after reactivating LXR signaling. It will be interesting to identify the LXR generated signals that revert the macrophage population in MASH and to see if these signals can be generated at other locations associated with altered myeloid populations including obese adipose tissue, atherosclerotic blood vessels and sites of amyloid beta deposition in Alzheimer’s disease.

Selectively expressing W441F in hepatocytes recapitulates the MASH phenotypes detected when the mutant protein is expressed from the endogenous locus except for the loss of Kupffer cells. Lipid accumulation in hepatocytes is thought to be the initial insult in the development of MASH ^2, 3, 66^. Our results suggest that the accumulation of hepatic cholesterol may be at least one critical trigger that drives subsequent inflammation and fibrosis. The cholesterol dependent phenotype seen in W441F mice is also consistent with studies in humans which correlate hepatic cholesterol levels with MASH ^5, 6, 8^. Cholesterol-dependent stabilization of the transcriptional cofactor TAZ in hepatocytes has been reported to contribute to the development of inflammation and fibrosis ^64^. Increased levels of TAZ protein are detected in the livers of male W441F mice after MASH diet feeding. Female W441F mice, however, have TAZ protein levels that are similar to controls. Taken together our results suggest that there may be TAZ-independent pathways that can promote MASH in response to high cholesterol levels.

Heterozygous W441F mice provide a unique example of the interplay between the environment in the form of diet and genotype. On a normal chow diet W/F mice exhibit decreased expression of LXR target genes, and consequently lower plasma lipid levels but otherwise appear normal. After 2 weeks of exposure to a MASH diet, however, these animals exhibit hepatic cholesterol accumulation, liver damage, inflammation, and fibrosis. There is also a corresponding change in the expression of over 4000 genes. Reactivation of LXR signaling in W/F mice reverses the effect of diet. Notably, the liver transcriptome in reactivated W/F mice closely resembles the transcriptome of vehicle treated control mice which are steatotic but show no evidence of MASH. Going forward, we suggest that W/F mice will provide a unique model system for identifying the factors that promote the development of MASH and for identifying potential therapeutic targets downstream of LXR signaling useful for reversing the disease.

## Methods

### Materials

Commercial reagents, chemicals, antibodies, mouse lines, cell lines, viruses, plasmids, and software used are listed in Supplemental Table 1. Sequence of the gRNA used to generate W441F mice and oligonucleotide sequences used for real-time PCR can be found in Supplemental Tabe 2.

### *In vitro* experiments

Immortalized bone marrow derived macrophage (BMDM) isolated from *Lxrα*^-/-^ + *Lxrβ*^-/-^ mice ^44^ (a gift from P. Tontonoz) were cultured in BMDM media (DMEM + 20% FBS + 30% L929 conditioned media) and infected with adenovirus expressing GFP, FLAG-LXRα or FLAG-LXRα W441F at a ratio of 100 infectious viral particles/cell for 24 hours. Following infection, cells were washed with phosphate buffered saline (PBS) and cultured for an additional 24 hours in BMDM media in the presence or absence of 10 µM 24,25-epoxycholesterol or 1.0 µM T0901317. After ligand treatment, RNA was isolated using Quick-RNA miniprep kits and mRNAs were quantified by real-time PCR. To examine the dominant negative activity of LXRα W441F, mouse AML12 cells cultured as recommended by the American Type Culture Collection were infected as described above. The media was changed 24 hours post-infection and RNA was isolated using Quick-RNA miniprep kits after an additional 24 hours. Real-time PCR was used to quantify mRNA levels. For Western blotting, 30 µg of nuclear extracts prepared as described by Liebergall et al. ^41^ were resolved on 10% SDS polyacrylamide gels, transferred to Immobilon-P membranes, probed with antibodies to FLAG, LXRα or TBP and visualized with the appropriate alkaline phosphatase conjugated second antibodies.

### Mice and *in vivo* experiments

The University of Virginia Animal care and Use Committee approved all animal experiments. CRISPR was used to replace tryptophan 441 at the Nr1h3 locus with phenylalanine. Fertilized eggs from BgSJLF1/J mice were injected with Cas9 protein, a gRNA targeting the region around tryptophan 441 and a 200 nucleotide replacement sequence that changes the TGG tryptophan codon at amino acid 441 to TTC encoding phenylalanine. This change also creates a novel TCGA sequence that is a site for cleavage by the restriction enzyme TaqI-v2. Founders were identified by a combination of restriction enzyme digestion and DNA sequencing. The W441F allele was then backcrossed 6 times into C57BL/6J. All experiments used offspring of heterozygous crosses. For experiments using the MASH diet, 11-12 week old male and female mice were fed Teklad Custom Diet TD.96121 ad libitum for 4 weeks. After 2 weeks mice were dosed daily by oral gavage with vehicle (80% polyethylene glycol 400/20% Tween 80) or with 10 mg/kg T0901317 for an additional 2 weeks. Mice were bled via the retro-orbital sinus and tissues were isolated approximately 3 hours after dosing. Body fat composition was measured using an EchoMRI^TM^ Whole Body Composition Analyzer (version 150521).

To generate hepatocyte-selective W441F mice, a cDNA encoding mouse LXRα W441F with a 7 amino acid FLAG tag at the amino terminus was cloned into the *Rosa26* targeting vector pR26 CAG AsiSI/MluI (a gift from Ralf Kuehn) downstream of a Lox-Stop-Lox cassette. Cas9 protein, the donor plasmid and a gRNA targeting *Rosa26* described by Chu et al. ^70^ were injected into fertilized eggs from BgSJLF1/J mice. Founders were identified by PCR and backcrossed 6 times with C57BL/6J mice. The final backcross utilized *Lxrα*^fl/fl^ mice, in the C57BL/6J background, that have previously been described ^23^. The offspring of the last backcross were bred to each other to produce *Rosa26*-LSL-FLAG-LXRα W441F + *Lxra*^fl/fl^ mice. To examine the response to the MASH diet, 9-10 week old female mice were injected via the tail vein with AAV-TBG-GFP or AAV-TBG-Cre (2×10^11^ viral genomes/mouse, gifts from James M. Wilson). After infection mice were maintained on a normal chow diet for 2 weeks before being switched to the MASH diet for 4 weeks.

### Plasma, hepatic, and fecal lipid measurements

Plasma total cholesterol, free cholesterol, triglycerides, glucose, ALT, and AST were measured using colorimetric assays listed in Supplemental Table 1 that were modified for use in 96 well plates. Hepatic lipids were extracted from approximately 50 mg of frozen samples as described by Breevoort et al. ^31^. Dried extracts were resuspended in isopropanol: Triton X 100 (9:1). Cholesterol and triglycerides were measured using the colorimetric assays described in Supplemental Table 1 and normalized to the amount of starting tissue. To measure fecal bile acids, mice were housed individually for 48 hours and feces from each animal was collected. Approximately 200 mg of feces was extracted as described by Breevoort et al. ^31^ and bile acids were measured using the colorimetric assay. Bile acids were normalized by the milligram amount of feces.

### Histochemistry and Immunostaining

Antibodies and other reagents for histochemical staining are described in Supplemental Table 1. Freshly harvested liver tissues were fixed in 4% formaldehyde for 48 hours and embedded in paraffin. For staining, 10 micron thick sections were deparaffinized, and stained with H&E or with Picrosirius Red (0.1% Direct Red 80 in a saturated aqueous solution of picric acid). To quantify collagen staining, ImagePro Plus was used to determine the percentage of Picrosirius Red positive area in 5 independent sections/mouse. For Oil Red O staining, freshly harvested liver and spleen samples were frozen in OTC and 10 micron thick sections were fixed in 60% ethanol for 10 minutes, stained with 0.21% Oil Red O in 60% isopropanol for 20 minutes, washed with water and then stained for 3 minutes with Hematoxylin QS. For immunostaining of Cd68, Clec4f, and Ki67, 10 micron thick frozen sections were fixed in 95% ethanol for 30 minutes and treated with BLOXALL for 10 minutes to inhibit endogenous peroxidase activity. Sections were then treated with Avidin and Biotin blocking solutions before incubation with primary antibodies for 1 hour followed by biotinylated second antibodies for 30 minutes. Staining was visualized using Avidin-Biotin Complex reagent followed by NovaRed staining. Nuclei were counter stained with Hematoxylin QS for 3 minutes.

### Liver Extracts and Western blotting

To prepare nuclear extracts approximately 50 mg of frozen liver in Lysing Matrix D tubes with 1.0 ml of Buffer A (10 mM HEPES pH 7.9, 10 mM KCl, 0.1 mM EDTA, 0.1 mM EGTA, 1.0 mM DTT, 0.6% NP-40, 1.0 mM PMSF, 25 µg/ml Aprotinin, 5 µg/ml Leupeptin, 1 µg/ml Pepstatin) were disrupted by sequential 30 second bursts in a FASTPREP 24 machine (MP Biochemicals) at speed setting 6 until a homogenous suspension was apparent. The suspension was then incubated on ice for an additional 5 minutes. Nuclei were pelleted for 1 minute at 16,000 x g at 4°C, the pellet was resuspended in 200 µl of Buffer A, pelleted as before and extracted in 50 µl of Buffer C (10 mM HEPES pH 7.9, 400 nM NaCl, 1.0 mM EDTA, 1.0 mM EGTA, 1.0 mM DTT, 0.6% NP-40, 1.0 mM PMSF, 25 µg/ml Aprotinin, 5 µg/ml Leupeptin, 1 µg/ml Pepstatin) for 15 minutes at 4°C with moderate shaking. Samples were pelleted for 5 minutes at 16,000 x g at 4°C, supernatants were collected, and protein amounts were quantified by BCA assays. For Western blotting, 30 µg of nuclear extract was resolved on 10% SDS polyacrylamide gels, transferred to Immobilon-P membranes, probed with antibodies to FLAG, LXRα or TBP and visualized with the appropriate alkaline phosphatase conjugated second antibodies. For whole cell extracts approximately 50 mg of frozen liver in Lysing Matrix D tubes with 0.5 ml of RIPA buffer (10 mM Tris pH 8.0, 1.0 mM EDTA, 1.0 mM EGTA, 0.1% SDS, 0.1% deoxycholate, 1.0% Triton X-100/1.0 mM PMSF, 25 µg/ml Aprotinin, 5 µg/ml Leupeptin, 1 µg/ml Pepstatin) was disrupted in a FASTPREP 24 Machine (MP Biochemicals) as described above. Extracts were incubated on ice for 5 minutes, pelleted for 10 minutes at 16,000 x g at 4°C, supernatants were collected, sheared with a 1cc Insulin syringe, and pelleted for 5 minutes at 16,000 x g at 4°C. Supernatants were collected and protein amounts were quantified by BCA assays. Western blotting was carried out as described above using antibodies recognizing TAZ and GAPDH.

### Liver macrophage isolation

Enriched populations of liver macrophages were isolated as described by Li et al. ^71^. Following isolation, red blood cells were lysed and the remaining cells were plated in 6 cm plates in 2.0 ml of RPMI media containing 10% fetal bovine serum and incubated at 37⁰C in 5% CO_2_ for 30 minutes. Cells were washed extensively with PBS and RNA was isolated using Quick-RNA miniprep kits.

### RNA Isolation from mouse tissues, Real-Time PCR, and RNA-Seq

Approximately 25 mg of tissue in Lysing Matrix D tubes with 0.5 ml of Triazole were disrupted for 30 second bursts in a FASTPREP 24 machine (MP Biochemicals) at speed setting 3. Following extraction with chloroform, total RNA was isolated from the aqueous layer using Quick-RNA miniprep kits. DNase treatment, reverse transcription and real-time PCR were carried out as described by Liebergall et al. ^41^ using a BIO-RAD CSF Connect system. Primers are listed in Supplemental Table 2. RNA-Seq using poly-A selected liver RNA isolated from male mice was carried out by Azenta Life Sciences. For each sample at least 20 million sequence reads were obtained. Sequence reads were trimmed to remove possible adapter sequences and nucleotides with poor quality using Trimmomatic v.0.36. The trimmed reads were mapped to the Mus musculus GRCm38 reference genome available on ENSEMBL using the STAR aligner v.2.5.2b. Unique gene hit counts were calculated by using featureCounts from the Subread package v.1.5.2. Only unique reads that fell within exon regions were counted. After extraction of gene hit counts, the gene hit counts table was used for downstream differential expression analysis. Using DESeq2, a comparison of gene expression between defined groups of samples was performed. The Wald test was used to generate p-values and log2 fold changes. Genes with an adjusted p-value < 0.05 and absolute log2 fold change > 1 were called as differentially expressed genes for each comparison.

### Statistical Analysis

All analysis was carried out using GraphPad Prism. Data sets were first tested for normal distributions. Experiments with normal distributions were examined using 1-way ANOVA, two-way ANOVA, or unpaired two-tailed t-tests. Two-tailed Mann-Whitney tests were used to data sets that did not meet a normal distribution.

## Acknowledgements

This work was supported by grants from the NIH/NIDDK (1R01DK130050-01A1) and NIH/NIA (1R21AGO75577-01A1) to I.G.S. Micrographs were obtained using the Leica Thunder Microscope in the University of Virginia Advanced Microscopy Facility. Mouse lines were made by the University of Virginia Genetically Engineered Mouse Model Core. Tissue sectioning and H&E staining was provided by the University of Virginia Research Histology Core. All core facilities are supported by the University of Virginia School of Medicine, and through the University of Virginia Cancer Center National Cancer Institute P30 Center Grant.

## Author Contributions

A.T.C, L.R-S., and I.G.S. designed experiments, performed experiments, and interpreted data. I.G.S. wrote the manuscript. L.A.A. and N.H. genotyped mice and performed histological staining.

## Materials and Correspondence

Correspondence and requests for materials should be made to Ira Schulman

## Data Availability

The authors declare that the data supporting the findings of this study are available within the paper and its supplementary information file and are available upon request from the corresponding author. Reagents and mouse lines are available upon request from the corresponding author. RNA sequencing data are available at the Gene Expression Omnibus accession number GSE267011.

