## Supplemental Figures and Tables for "A Novel Mutation in LXRα Uncovers a Role for Cholesterol Sensing in Limiting Metabolic Dysfunction-Associated Steatohepatitis (MASH)"

### Supplemental Figure 1

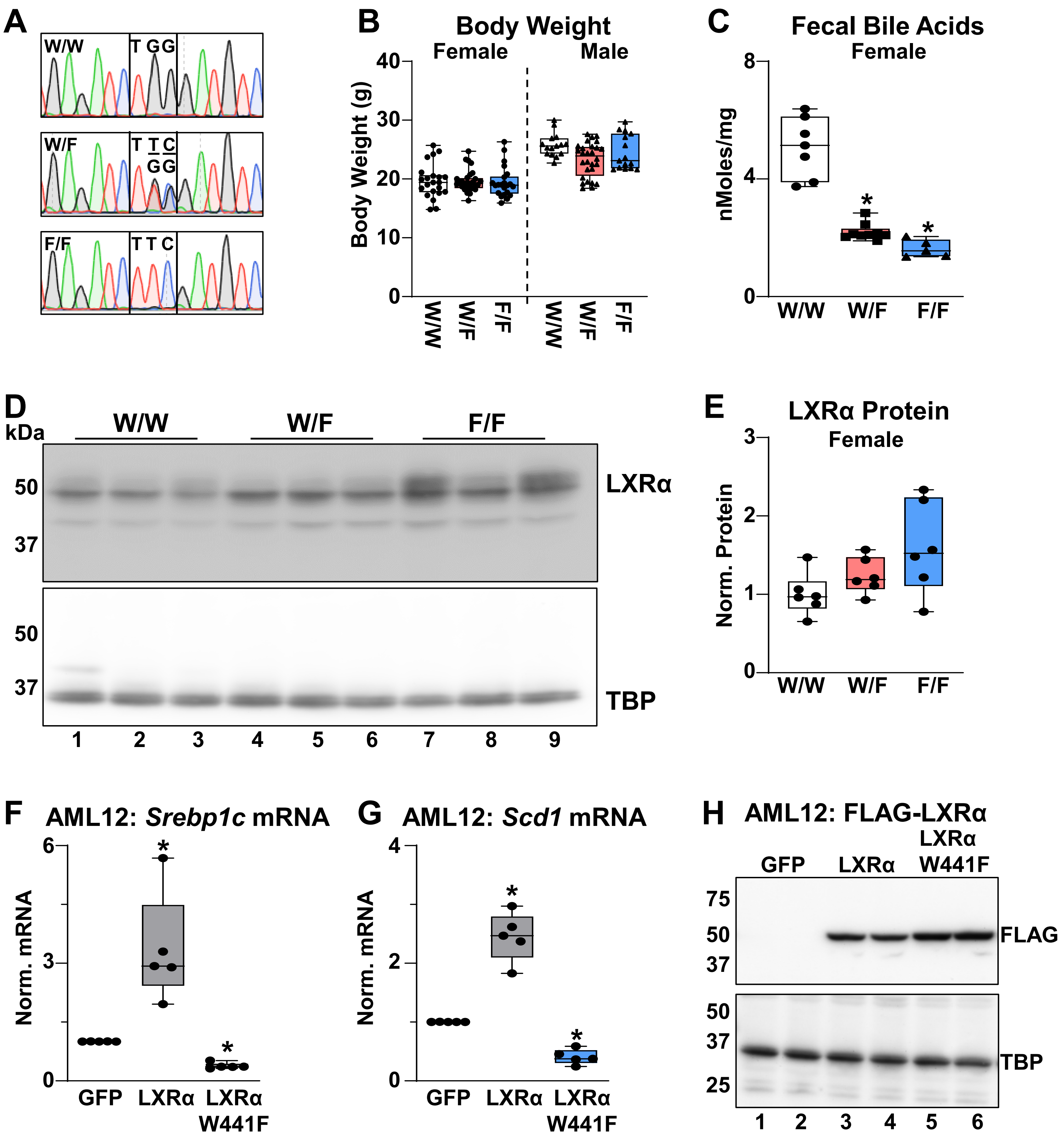

**Supplemental Figure 2**

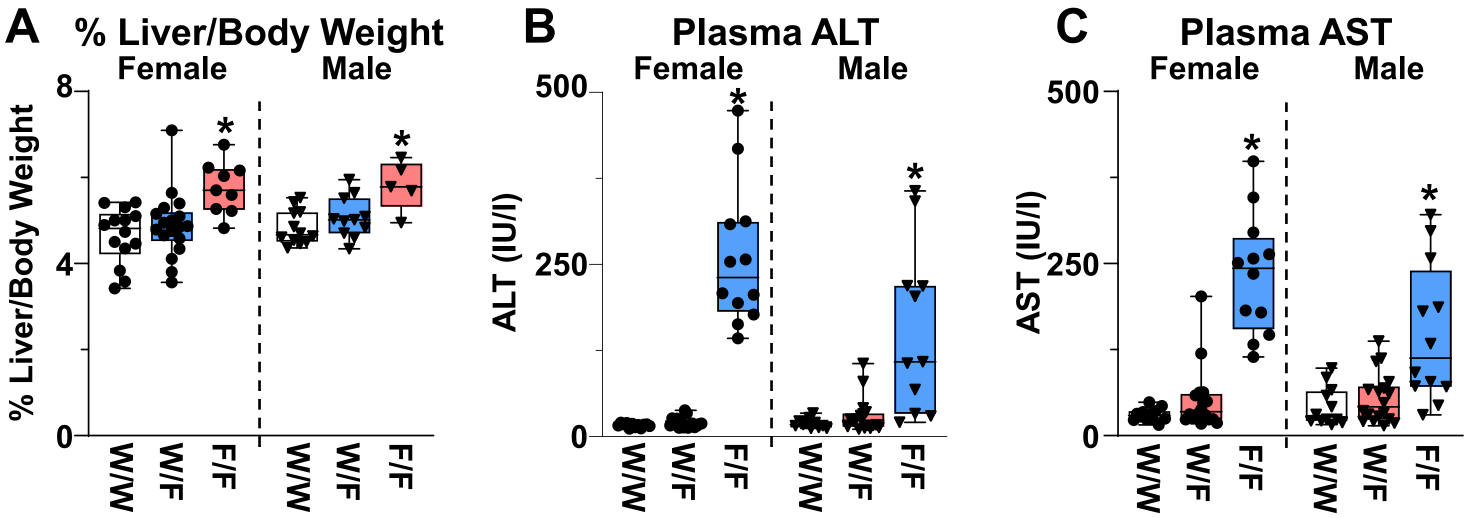

#### Supplemental Figure 3

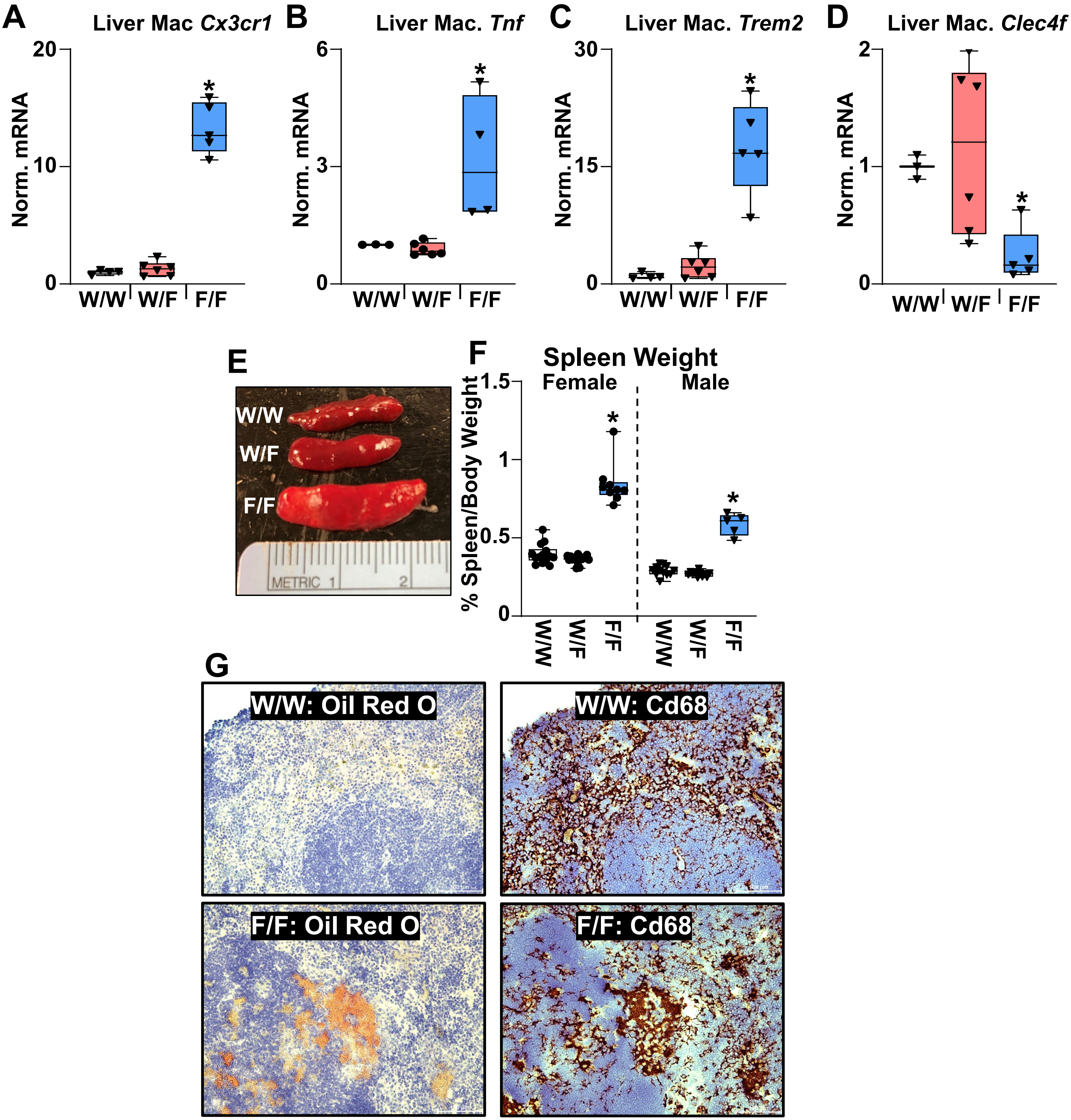

Supplemental Figure 4

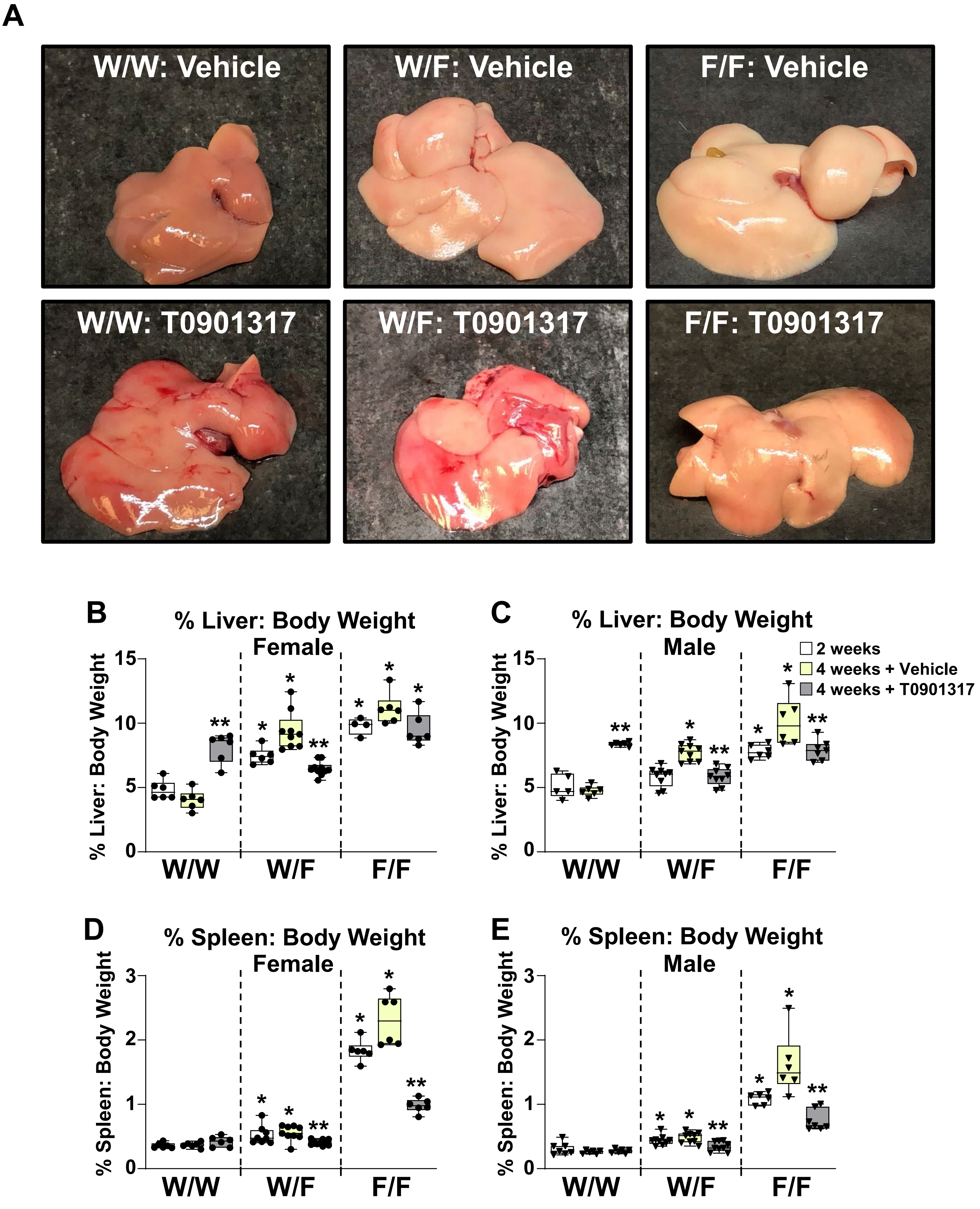

Supplemental Figure 5

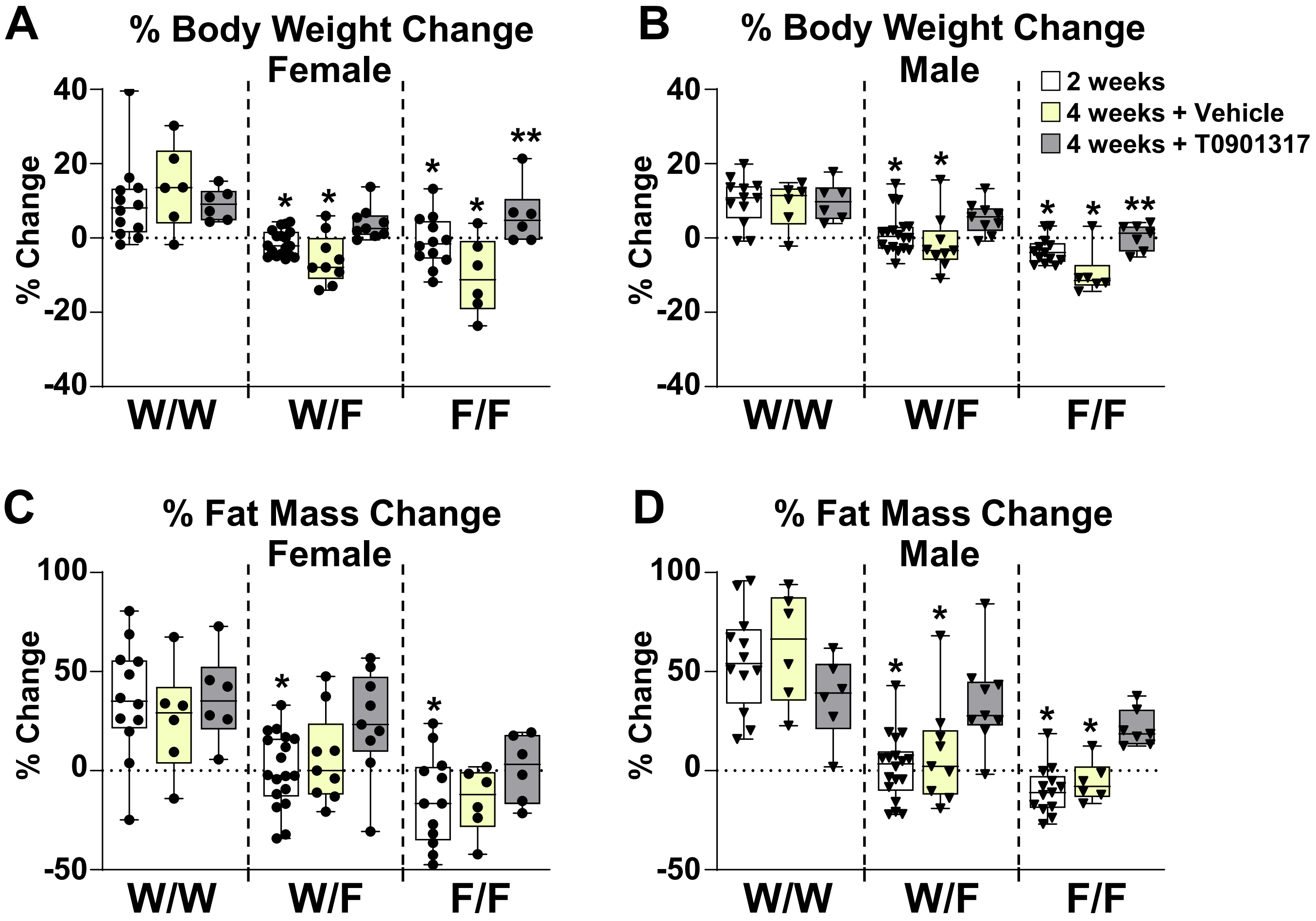

#### Supplemental Figure 6

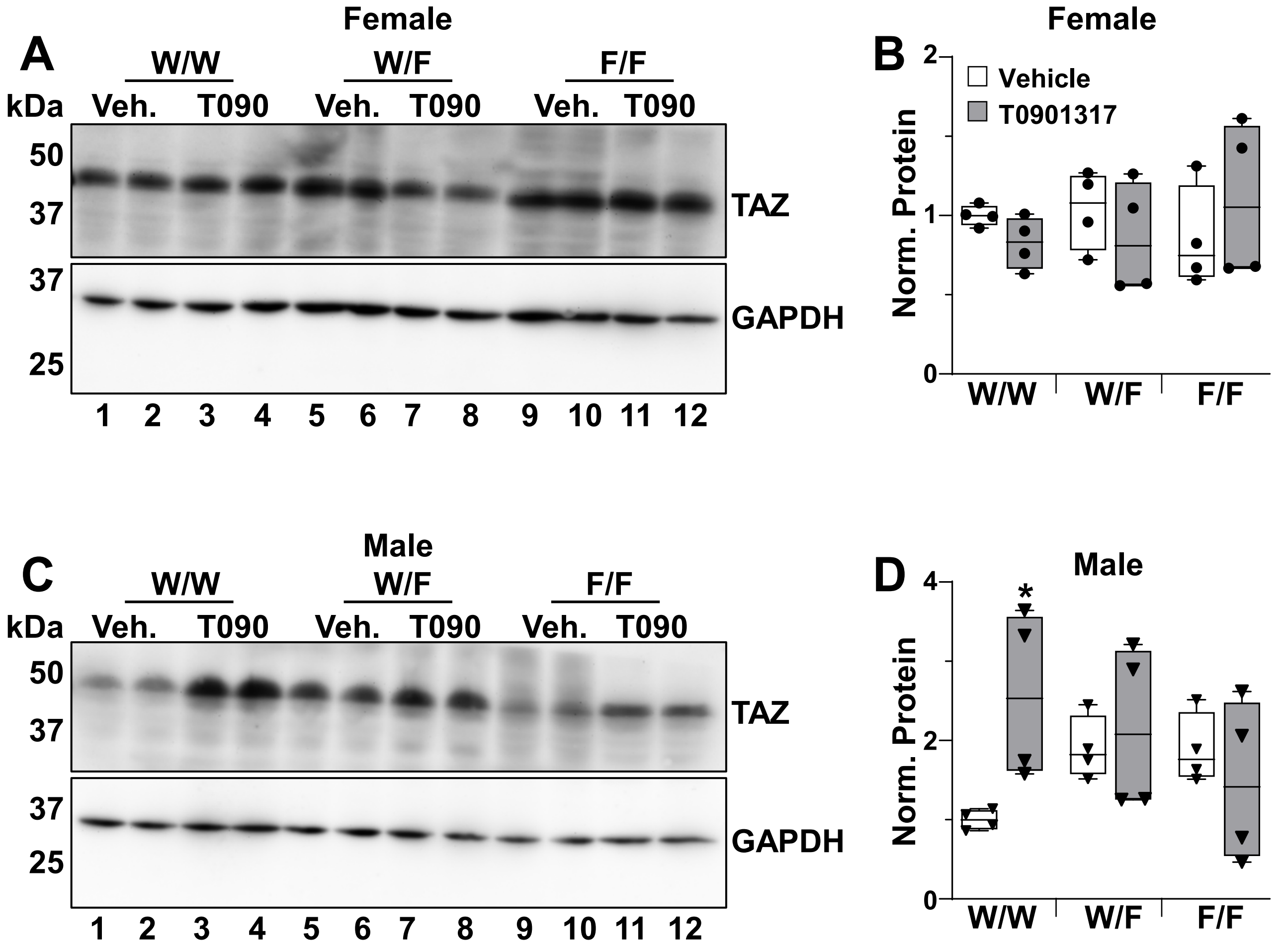

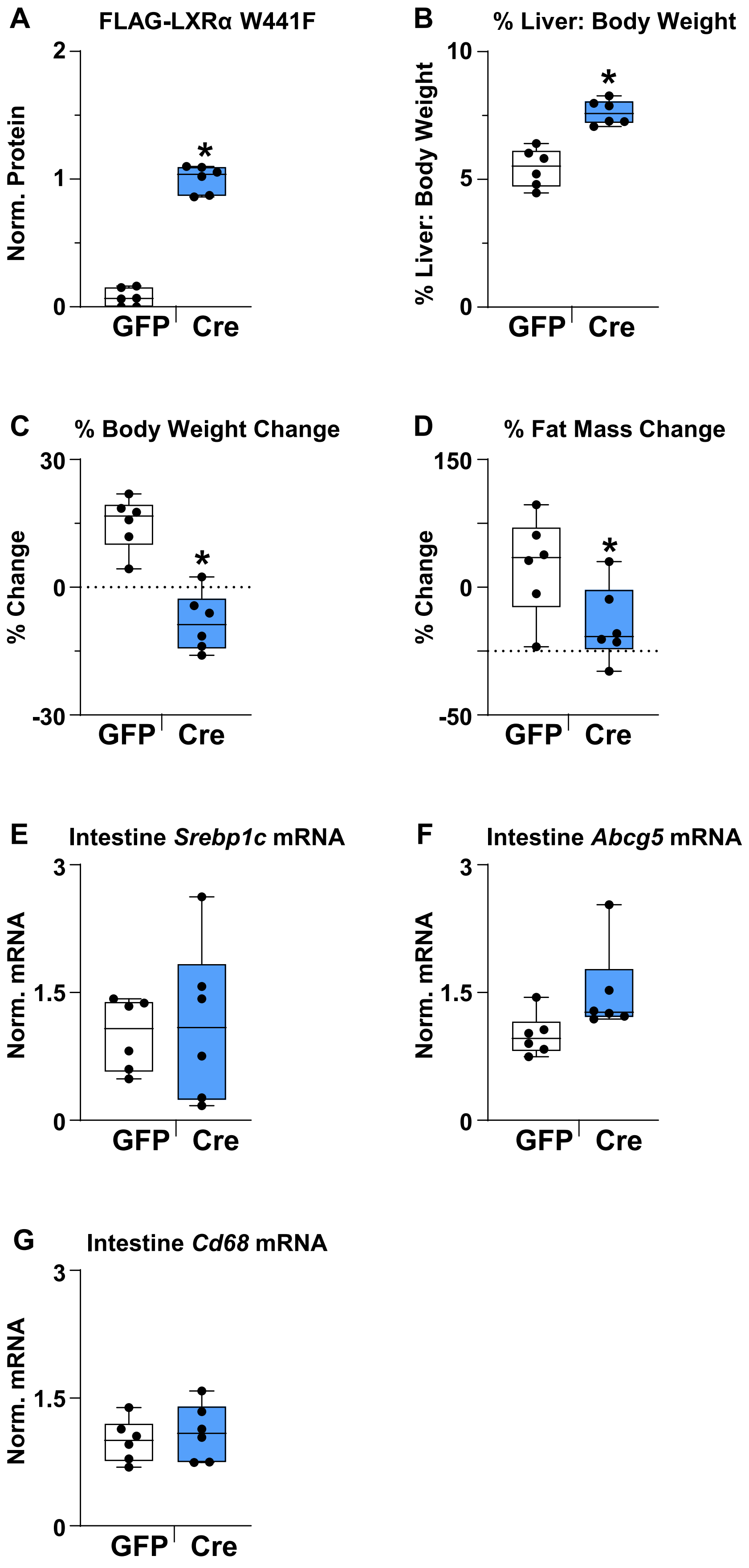
A
FLAG-LXRα W441F
Norm. Protein
2
1
0
*
GFP
Cre
B
% Liver: Body Weight
% Liver: Body Weight
10
5
0
*
GFP
Cre
C
% Body Weight Change
% Change
30
0
-30
*
GFP
Cre
D
% Fat Mass Change
% Change
150
0
-50
*
GFP
Cre
E
Intestine *Srebp1c* mRNA
Norm. mRNA
3
1.5
0
GFP
Cre
F
Intestine *Abcg5* mRNA
Norm. mRNA
3
1.5
0
GFP
Cre
G
Intestine *Cd68* mRNA
Norm. mRNA
3
1.5
0
GFP
Cre

#### Supplemental Figure Legends

**Supplemental Figure 1. Characterization of LXRα W441F mice. A**) DNA Sequence of amino acid 441 illustrating the change from tryptophan (TGG) to phenylalanine (TTC). **B**) Body weight of 11-12 week old control (W/W), heterozygous mutant (W/F), and homozygous mutant (F/F) W441F mice maintained on normal chow diet (n = 15-28/group). **C**) Fecal bile acids were measured in 11-12 week old female mice as described in the Methods section. *Statistically significant difference between control and mutant mice determined by one-way ANOVA ($p \leq 0.05$, n = 5-7/group). **D**) Liver nuclear extracts were prepared as described in the Methods section. LXRα and TBP were examined by Western blotting. Each lane represents an individual mouse. **E**) LXRα protein levels in female mice were quantified from Western blots by normalization to TBP levels using ImageQuant TL (n = 6/group). Each point is an individual mouse. The amount of LXRα in control W/W mice was set at 1. **F**-**H**) AML12 cells were infected with adenovirus expressing LXRα or LXRα W441F. RNA was harvested 48 hours after infection and the mRNA levels of **F**) *Srebp1c* and **G**) *Scd1* were measured by real-time PCR as described in the Methods section. *Statistically significant difference between GFP and LXRs determined by one-way ANOVA ($p \leq 0.05$, n =5). The line in the middle of the bars represents the median. **H**) Nuclear extracts were prepared as described in the Methods section. FLAG-LXRα and TBP levels were examined by Western blotting.

**Supplemental Figure 2. Liver weight and liver enzymes in LXRα W441F mice**. **A**) Percent liver to body weight ratio in female and male 11-12 week old control (W/W), heterozygous mutant (W/F), and homozygous mutant (F/F) W441F mice maintained on normal chow diet. *Statistically significant difference between W/W and mutant mice determined by one-way

ANOVA ($p \leq 0.05$, n = 5-12/group). **B**) Plasma ALT and **C**) AST levels in female and male 11-12 week old control (W/W), heterozygous mutant (W/F), and homozygous mutant (F/F) W441F mice maintained on normal chow diet. *Statistically significant difference between control W/W and mutant mice determined by one-way ANOVA ($p \leq 0.05$, n = 11-18/group). The line in the middle of the bars represents the median.

**Supplemental Figure 3. Liver and splenic macrophages from LXRα W441F mice**. **A-D**) Gene expression measured by real-time PCR from enriched populations of liver macrophage isolated from 8-10 week old male control (W/W), heterozygous mutant (W/F), and homozygous mutant (F/F) W441F mice maintained on normal chow diet as described in the Methods section. *Statistically significant difference between control W/W and mutant mice determined by one-way ANOVA ($p \leq 0.05$, n = 3-6/group). **E**) Representative spleens from 12 week old male control (W/W), heterozygous mutant (W/F), and homozygous mutant (F/F) W441F mice maintained on normal chow diet. **F**) % spleen weight to body weight ratio from 11-12 week old control (W/W), heterozygous mutant (W/F), and homozygous mutant (F/F) W441F mice maintained on normal chow diet. *Statistically significant difference between control W/W and mutant mice determined by one-way ANOVA ($p \leq 0.05$, n = 5-18/group). The line in the middle of the bars represents the median. **G**) Spleen sections from 12 week old male control (W/W) and homozygous mutant (F/F) W441F mice stained with Oil Red O and antibodies to Cd68 as described in the Methods sections.

**Supplemental Figure 4. Liver and spleen weights from LXRα W441F mice fed the MASH diet**. Female and male 11-12 week old control (W/W), heterozygous mutant (W/F), and homozygous mutant (F/F) W441F mice were fed the MASH diet. A subset of mice was sacrificed after 2 weeks. The remaining mice were maintained on the MASH diet for an additional 2 weeks and treated with vehicle or 10 mg/kg T0901317 daily by oral gavage as described in the Methods section. **A**) Representative livers from male mice treated with vehicle or T01901317 for the final 2 weeks of the 4 week MASH diet feeding experiment. **B**-**C**) Percent liver weight to body weight. **D**-**E**) Percent spleen weight to body weight. *Statistically significant difference between control and mutant mice in the same treatment group; **statistically significant difference between vehicle and T0901317 treated mice of the same genotype determined by two-way ANOVA ($p \leq 0.05$, n = 5-9/group). The line in the middle of the bars represents the median.

**Supplemental Figure 5. Body weight and fat mass changes in LXRα W441F mice fed the MASH diet**. Female and male 11-12 week old control (W/W), heterozygous mutant (W/F), and homozygous mutant (F/F) W441F mice were fed the MASH diet. A subset of mice was sacrificed after 2 weeks. The remaining mice were maintained on the MASH diet for an additional 2 weeks and treated with vehicle or 10 mg/kg T0901317 daily by oral gavage as described in the Methods section. **A**-**B**) Percent body weight and **C**-**D**) percent fat mass change relative to day 1 measured by Echo/MRI. *Statistically significant difference between control and mutant mice in the same treatment group; **statistically significant difference between vehicle and T0901317 treated mice of the same genotype determined by two-way ANOVA ($p \leq 0.05$, n = 6-18/group). The line in the middle of the bars represents the median.

**Supplemental Figure 6. TAZ protein levels in LXRα W441F mice fed the MASH diet**. Female and male 11-12 week old control (W/W), heterozygous mutant (W/F), and homozygous mutant (F/F) W441F mice were fed the MASH diet for 4 weeks and treated with vehicle or 10 mg/kg T0901317 daily by oral gavage for the final 2 weeks. After drug treatment, liver whole cell extracts were prepared and levels of TAZ and GAPDH were examined by Western blotting (**A** and **C**). Each lane represents an individual mouse. TAZ protein levels (**B** and **D**) were quantified from Western blots by normalization to GAPDH using ImageQuant Software (n = 4/group). TAZ levels in vehicle treated control W/W mice were set at 1. *Statistically significant difference between vehicle and T0901317 treated mice of the same genotype by two-way ANOVA ($p \leq 0.05$, n = 4/group). The line in the middle of the bars represents the median.

**Supplemental Figure 7. Characterization of *Rosa26*-LSL-FLAG-LXRα W441F/*Lxra*$^{fl/fl}$ mice**. Female 9 week old *Rosa26*-LSL-FLAG-LXRα W441F/*Lxra*$^{fl/fl}$ mice were infected with AAV-TBG-GFP or AAV-TBG-Cre viruses. After infection, mice were maintained on a normal chow diet for 2 weeks and then fed the MASH diet for 4 weeks. **A**) FLAG-LXRα W441F protein levels in liver nuclear extracts were quantified from Western blots by normalization to TBP using ImageQuant Software (n = 6/group). FLAG-LXRα W441F levels in Cre mice were set as 1. **B**) Percent liver weight to body weight ratio, **C**) percent body weight change, and **D**) percent fat mass change relative to the start of the MASH diet. *Statistically significant difference between GFP and Cre mice determined by unpaired two-tailed t-test ($p \leq 0.05$, n = 6/group). **E-G**) Intestine gene expression measured by real-time PCR as described in the Methods section. *Statistically significant difference between GFP and Cre mice determined by unpaired two-tailed t-test or two-tailed Mann-Whitney test ($p \leq 0.05$, n = 6/group). The line in the middle of the bars represents the median.

**Supplemental Table 1. Materials and Reagents**

| Commercial Reagents and Chemicals | Source | Identifier |
|---|---|---|
| Total Cholesterol Assay | Fisher Scientific | TR13421 |
| Free Cholesterol Assay | FujiFilm/Wako | 993-02501 |
| Triglycerides Assay | Fisher Scientific | T7532-120 |
| Glucose Assay | Fisher Scientific | G7517-120 |
| ALT Assay | Fisher Scientific | A7526-150 |
| AST Assay | Fisher Scientific | A7561-150 |
| Bile Acids Assay | Fisher Scientific | DZ042A-KY1 |
| MASH Diet | Teklad | TD.96121 |
| 24,25-epoxycholesterol | Cayman Chemicals | 10131 |
| T0901317 | Cayman Chemicals | 71810 |
| Triazole | Fisher Scientific | 15596026 |
| BCA Asay Kit | Fisher Scientific | 23225 |
| Direct Red | MilliporeSigma | 365548-5G |
| Oil Red O | MilliporeSigma | O0625-25G |
| BLOXALL | Vector Laboratories | SP-6000 |
| Avidin,/Biotin Blocking Kit | Vector Laboratories | SP-2001 |
| Vectastain Elite ABC Reagent, Peroxidase | Vector Laboratories | SK-4800 |
| Vector NovaRed Substrate kit, Peroxidase | Vector Laboratories | SK-4800 |
| Hematoxylin QS | Vector Laboratories | H-3404 |
| Lysing Matrix D Tubes | MP Biomedicals | 6913500 |
| Quick-RNA Miniprep Kits | Zymo Research | R1055 |
| Immobilon-P membrane | MilliporeSigma | IPVD00010 |
| **Antibody (dilution)** | **Source** | **Identifier** |
| Anti-Cd68 (1/1500) | Biolegend | 137001 |
| Anti-Clec4f (1/1000) | R&D Systems | 2785-CL |
| Anti-GAPDH (1/1000) | Cell Signaling | 5174S |
| Anti-Ki67 (1/500) | Cell Signaling | 12202T |
| Anti-LXRα (1/3000) | R&D Systems | PP-PPZ0412-00 |
| Anti-TAZ (1/1000) | R&D Systems | 83669S |
| Anti-TBP (1/1000) | Cell Signaling | 440595 |
| Goat Anti-Mouse IgG+IgM Alkaline Phosphatase (1/5000) | Fisher Scientific | T2192 |
| Goat Anti-Rabbit IgG+IgM Alkaline Phosphatase (1/5000) | Fisher Scientific | T2191 |
| Goat Anti-Rabbit IgG Biotinylated (1/200) | Vector Laboratories | BA-1000 |
| Rabbit Anti-Goat IgG Biotinylated (1/200) | Vector Laboratories | BA-5000 |
| Rabbit Ant-Rat IgG Biotinylated (1/200) | Vector Laboratories | BA-4001 |
| **Experimental Models: Organisms/Strains** | **Source** | **Identifier** |
| Mouse: BgSJLF1/J | Jackson Laboratories | 100012 |
| Mouse: C57BL/6J | Jackson Laboratories | 000664 |
| Mouse: *Lxrα*$^{fl/fl}$ | Mangelsdorf Lab | (Zhang et al., 2012) |

| | | |
|---|---|---|
| Mouse: *Lxrα* W441F | University of Virginia | This study |
| Mouse: *Rosa26* FLAG-LXRα W441F/ *Lxrα*$^{fl/fl}$ | University of Virginia | This study |
| Mouse: *Lxrα*$^{-/-}$/*Lxrβ*$^{-/-}$ Immortalized BMDM | Tontonoz Lab | (Ito et al., 2015) |
| Mouse: AML12 Cells | ATCC | CRL-2254 |
| **Virus and Plasmids** | **Source** | **Identifier** |
| AAV-TBG-Cre | Addgene | 107787-AAV8 |
| AAV-TBG-GFP | Addgene | 105535-AAV8 |
| pR26 CAG AsiSI/MluI | Addgene | 74286 |
| pAV-CMV-FLAG-mLXRα-PGK-EGFP | VectorBuilder | VB200409-1065yuy |
| pAV-CMV-FLAG-mLXRα W441F-PGK-EGFP | VectorBuilder | VB220906-1310evw |
| pAV-CMV-EGFP | Vector Builder | VB010000-9299haqc |
| **Software** | **Source** | |
| GraphPad Prism 10 | GraphPad Software | |
| Image-Pro Plus 7.0 | Media Cybernetics | |
| ImageQuant TL | Cytiva Life Sciences | |

**Supplementary Table 2. Oligonucleotides for CRISPR and qPCR.**

| Species | Gene | Sequence |
|---|---|---|
| Mouse | gRNA for W441F | 5’ CCCCTCTGCTGTCTGAGATCTGG 3’ |
| Mouse | Abca1 | 5’GCTCTCAGGTGGGATGCAG 3’<br>5’GGCTCGTCCAGAATGACAAC 3’ |
| Mouse | Abcg5 | 5’TGGATCCAACACCTCTATGCTAAA 3’<br>5’ GGCAGGTTTTCTCGATGAACTG 3’ |
| Mouse | Ccna2 | 5’GCCTTCACCATTCATGTGGAT 3’<br>5’CTTGCAGTGAGTGACGTAGAC 3’ |
| Mouse | Cd68 | 5’TTGGGAACTACACACGTGGGC 3’<br>5’CGGATTTGAATTTGGGCTTG 3’ |
| Mouse | Cdk1 | 5’AGAAGGTACTTACGGTGTGGT 3’<br>5’GAGAGATTTCCCGAATTGCAGT 3’ |
| Mouse | Clec4f | 5’GAGGCCGAGCTGAACAGAG 3’<br>5’TGTGAAGCCACCACAAAAAGAG 3’ |
| Mouse | Col1a1 | 5’GAGCAGACGGGAGTTTCTCCT 3’<br>5’CATGTAGACTCTTTGCGGCTG3’ |
| Mouse | Cyp7a1 | 5’GCTTGTAGAGAGCCACACCAA 3’<br>5’AGTGGTGGCAAAATTCCCA 3’ |
| Mouse | Cyp27a1 | 5’CCAGGCACAGGAGAGTACG 3’<br>5’GGGCAAGTGCAGCACATAG 3’ |
| Mouse | Cx3cr1 | 5’GAGTATGACGATTCTGCTGAGG 3’<br>5’CAGACCGAACGTGAAGACGAG 3’ |
| Mouse | Fasn | 5’CGGAAACTTCAGGAAATGTCC 3’<br>5’TCAGAGACGTGTCACTCCTGG 3’ |
| Mouse | Ppia (Cyclophilin) | 5’CGATGACGAGCCCTTGG 3’<br>5’TCTGCTGTCTTTGGAACTTTGTC 3’ |
| Mouse | Scd1 | 5’CCGGAGACCCCTTAGATCGA 3’<br>5’TAGCCTGTAAAAGATTTCTGCAAACC 3’ |
| Mouse | Spp1 | 5’ATCTCACCATTCGGATGAGCTT 3’<br>5’ TGTAGGGACGATTGGAGTGAAA 3’ |
| Mouse | Srebf1c | 5’CGTCTGCACGCCCTAGG 3’<br>5’CTGGAGCATGTCTTCAAATGTG 3’ |
| Mouse | Tgfb1 | 5’CTCCCGTGGCTTCTAGTGC 3’<br>5’ GCCTTAGTTTGGACAGGATCTG 3’ |
| Mouse | Tnf | 5’CTGAGGTCAATCTGCCCAAGTAC 3’<br>5’CTTCACAGAGCAATGACTCCAAAG 3’ |
| Mouse | Trem2 | 5’CTGGAACCGTCACCATCACTC 3’<br>5’CGAAACTCGATGACTCCTCGG 3’ |
